# Lung [^18^F]FDG-PET is an early pharmacodynamic biomarker of antiviral efficacy in preclinical influenza

**DOI:** 10.64898/2026.10.06.756978

**Authors:** Carla Bianca Luena Victorio, Arun Ganasarajah, Shantanu Gupta, Joanne Ong, Ann-Marie Chacko

## Abstract

**Background:** Influenza remains a major cause of global morbidity and mortality. Pharmacodynamic biomarkers that quantify treatment response within the lung could accelerate influenza therapeutic development. We evaluated [¹ F]fluorodeoxyglucose positron emission tomography ([^18^F]FDG-PET) as a non-invasive biomarker of antiviral efficacy.

**Methods:** Serial [^18^F]FDG-PET was evaluated in two murine H1N1 influenza models using mouse-adapted PR8 strain and CA-09 clinical isolate following treatment with favipiravir, baloxavir marboxil, or ribavirin. PET measures were compared with pulmonary viral burden, inflammatory cytokine expression, computed tomography (CT) imaging, and respiratory function measured by whole-body plethysmography (WBP).

**Results:** Antiviral treatment in PR8 infection model reduced lung [^18^F]FDG uptake by 27–49% and metabolic lung volume (MLV) by 40.5–74.6% (both *p* < 0.001). Antiviral response was detectable as early as 3 days post-infection, when uptake decreased by 28.7–53.2% and MLV by 33.8–75.0%. Lung [^18^F]FDG uptake also strongly correlated with MLV (*r* = 0.93, *p* < 0.001) and with pulmonary viral burden and inflammatory mediators. These findings were reproduced in a model of pandemic H1N1/CA-09, where baloxavir reduced uptake by 37.4% (*p* = 0.02) and MLV by 67.5% (*p* < 0.001) at day 3. In both models, changes in [^18^F]FDG-PET parameters preceded improvements in lung CT and respiratory function.

**Conclusions:** Lung [^18^F]FDG-PET provides an early, quantitative measure of antiviral response in experimental influenza, supporting its further evaluation as a translational pharmacodynamic endpoint.

## Background

Influenza remains a major cause of morbidity and mortality worldwide, resulting in an estimated 290,000-650,000 respiratory deaths and 3-5 million severe cases annually [1–3]. Beyond acute respiratory disease, influenza is associated with cardiovascular, renal, and neurological complications [4–7], with particularly severe outcomes in individuals with chronic conditions such as diabetes, asthma, and chronic obstructive pulmonary disease (COPD) [8, 9]. Persistent impairment in lung function and reduced quality of life after infection further contributes to disease burden [10, 11].

Seasonal vaccination and antiviral therapy remain the main strategies for influenza prevention and treatment. Neuraminidase inhibitors and polymerase inhibitors (favipiravir and baloxavir) reduce disease severity, while baloxavir can decrease household transmission [12–15]. However, antiviral resistance [16] and recurrent vaccine mismatch [17, 18] continue to drive development of antivirals with novel mechanisms, combination therapies, and broadly protective vaccines [19]. A major barrier to evaluating these interventions is the lack of biomarkers that directly and longitudinally quantify treatment effects within the lung, the principal site of severe influenza pathology. Circulating inflammatory proteins and breath-derived metabolites provide measures of host response [20–22] but cannot define the spatial extent or evolution of pulmonary inflammation. A non-invasive biomarker capable of repeatedly measuring lung inflammation *in vivo* could therefore provide an important intermediate measure of therapeutic response between virological endpoints and conventional clinical or physiological outcomes.

Positron emission tomography (PET) with [^18^F]fluorodeoxyglucose ([^18^F]FDG) provides spatially resolved quantification of inflammatory metabolic activity *in vivo* [23, 24]. [^18^F]FDG-PET has demonstrated utility as a pharmacodynamic readout in tuberculosis [25] and as inflammation biomarker across viral disease models [26–29], including respiratory viral infections MERS-CoV and SARS-CoV [30, 31]. Moreover, pulmonary [^18^F]FDG uptake in influenza correlates with disease severity and lung inflammation [32, 33]. Unlike systemic biomarkers or terminal tissue measurements, serial PET can quantify the magnitude and distribution of pulmonary inflammatory activity within the same subject over the course of disease and treatment. This creates the possibility of directly measuring whether an antiviral intervention modifies lung pathology, rather than relying solely on viral burden or downstream systemic responses.

We previously identified pulmonary [^18^F]FDG uptake as a sensitive marker of influenza-associated lung inflammation that correlated with viral replication and inflammatory responses in severe H1N1 infection [33]. Whether this imaging signal is responsive to effective antiviral therapy, and therefore suitable as a pharmacodynamic biomarker, remains to be established. Here, we address this question in two murine influenza models using mouse-adapted H1N1 A/Puerto Rico/8/34 (PR8) strain and clinical isolate H1N1 A/California/04/09 (CA-09). We evaluate whether serial [^18^F]FDG-PET detects antiviral-mediated suppression of lung inflammation and examine its relationship with pulmonary viral burden, inflammatory cytokines, and respiratory function measured by whole-body plethysmography (WBP). This study tests the potential of treatment-responsive pulmonary imaging as a translational pharmacodynamic endpoint for preclinical evaluation of influenza therapeutics.

## Results

### Lung [^18^F]FDG-PET detects early antiviral responses preceding clinical and structural disease resolution

To profile the pharmacodynamic sensitivity of lung [^18^F]FDG uptake, mice infected with H1N1 PR8 were treated with favipiravir (FPV), ribavirin (RBV), baloxavir (BXV), or vehicle prior to [^18^F]FDG-PET scans (**Fig. 1a**). Vehicle-treated mice developed severe disease and succumbed early (median survival, OS_m_ = 6 days), whereas all antivirals prevented mortality (OS_m_ >28 days; *p* < 0.001 for FPV, RBV, and BXV) (**Fig. 1b, c**) and improved clinical outcomes (**Fig. S1**). Antiviral treatment also reduced lung viral burden by 38–44% at peak disease and decreased viral antigen staining and lung pathology (**Fig. 1c,d**).

**Fig. 1.**
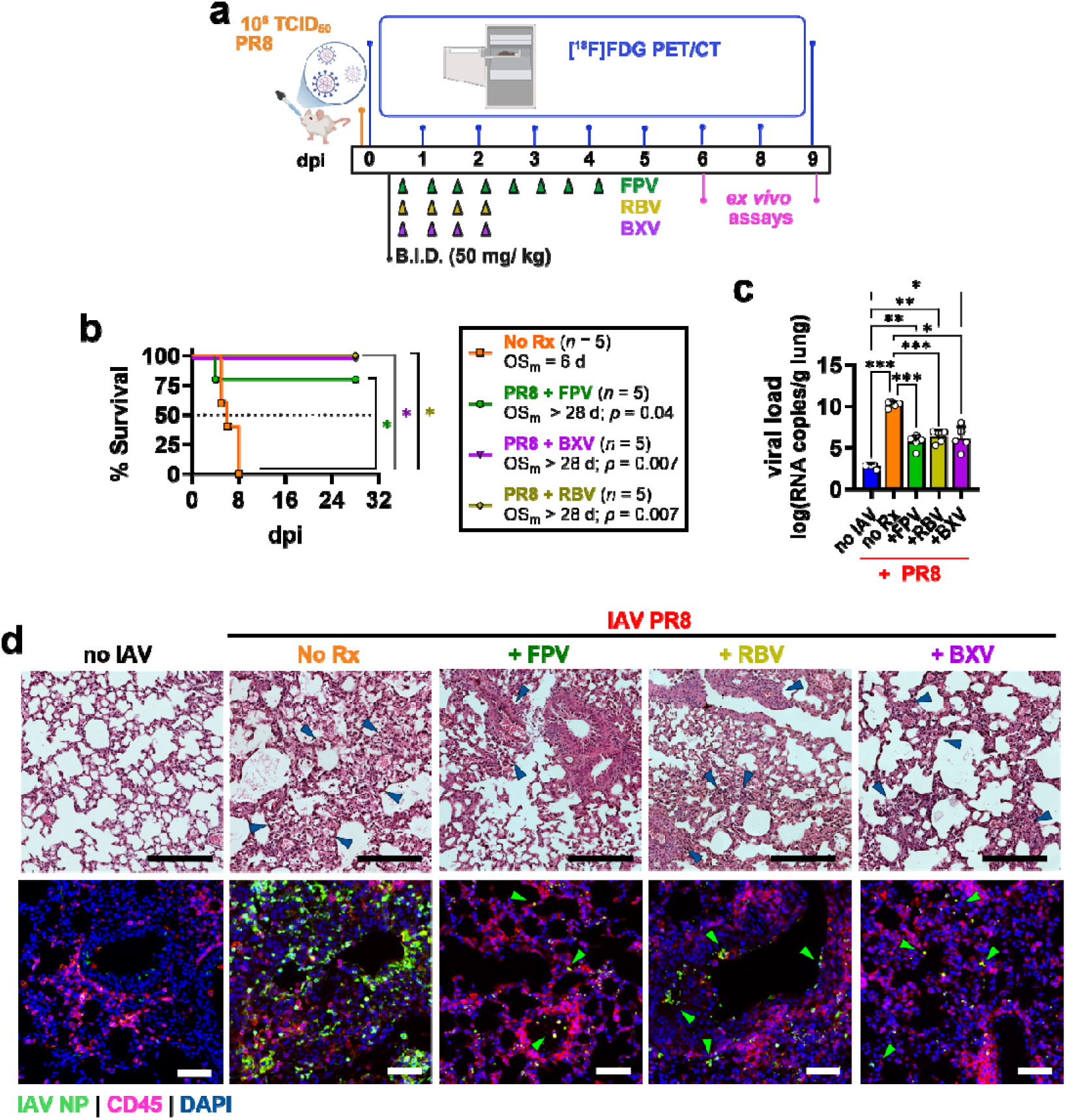
Mitigation of disease severity following antiviral therapy in a murine model of influenza A/PR8 infection. **a,** Study design depicting virus inoculation, antiviral treatment schedule, imaging timepoints, and endpoint *ex vivo* assays. Mice were infected with virus and treated twice daily with favipiravir (FPV), ribavirin (RBV), or baloxavir marboxil (BXV); or vehicle control (No Rx). Serial [¹ F]FDG-PET/CT imaging was performed from 0–6 days post-infection (dpi) in untreated mice and extended to 9 dpi in treated groups. *Ex vivo* analyses were performed at study endpoints. **b,** Kaplan– Meier survival curves. Dotted line indicates 50% survival. **c,** Lung viral load measured by quantitative RT–PCR at 6 dpi (untreated) or 9 dpi (treated). **d,** Representative lung immunohistopathology images on lung tissue sections (5 µm). H&E staining showing inflammatory cell infiltration (*blue arrows*) and immunofluorescence staining showing influenza A virus nucleoprotein (NP; *green arrows*) and CD45^+^ immune cell infiltration. Survival curves in ***b*** were compared with Mantel-Cox test with Holm-Šídák correction. Data in ***c*** reported as mean ± s.d from 5 mice per group were compared using Welch’s ANOVA with Dunnett’s correction. * *p* < 0.05; ** *p* < 0.005; ** *p* < 0.001; *ns*, not significant. Scale bars, 200 μm.

Serial [^18^F]FDG-PET revealed progressively increasing lung uptake in vehicle-treated mice through 6 days post-infection (dpi), whereas all antivirals suppressed this increase (**Fig. 2a,b**). PET imaging visualized progression of pulmonary inflammation and bone marrow hyperplasia in untreated animals and its attenuation following therapy (**Fig. 2c,d**). In contrast, lung CT did not distinguish treatment groups before 6 dpi (**Fig. S2**).

**Fig. 2.**
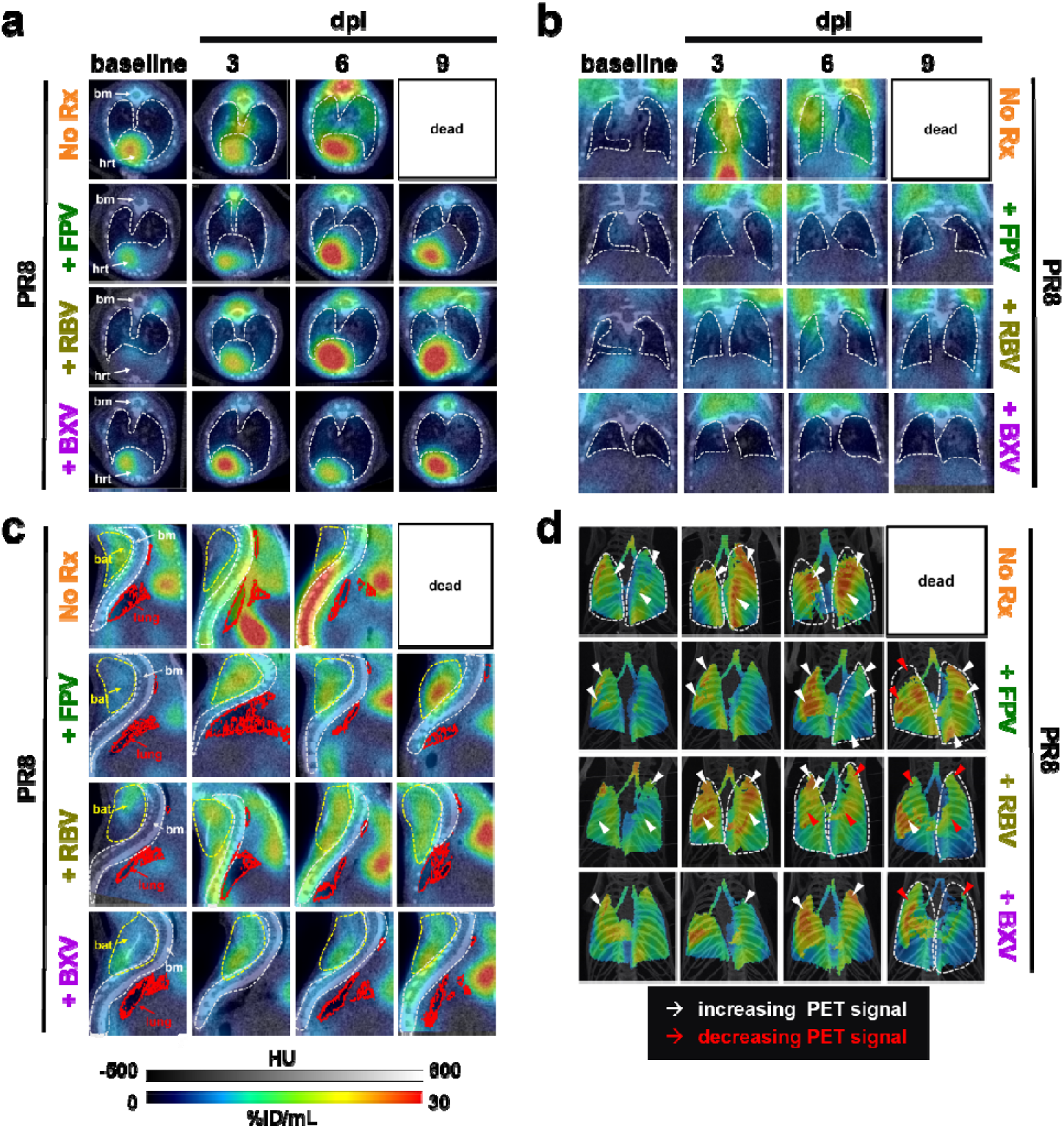
Dynamics of lung metabolism revealed by [¹ F]FDG-PET following antiviral treatment in PR8-infected mice. **a–c,** Representative [^18^F]FDG-PET/CT image slices shown in **a**, coronal; **b**, transverse; and **c**, sagittal views of lungs from influenza infected mice treated with either vehicle control (no Rx) or with antivirals favipiravir (FPV), ribavirin (RBV), or baloxavir marboxil (BXV), on various days post-infection (dpi). **d,** Representative maximum intensity projection (MIP) images showing temporal changes in lung [¹ F]FDG uptake. White arrows depict temporal increase in signal; while red arrows depict temporal decrease. Lung volumes of interest (VOIs) are indicated by grey dashed lines in ***a***, ***b***, and ***d***; and by red lines in ***c.*** CT images are displayed at −500 to 600 Hounsfield units (HU), and PET images at 0–30% ID/mL. *hrt,* heart; *bat,* brown adipose tissue; *bm,* bone marrow.

Over 6 days, antiviral treatment reduced lung [^18^F]FDG uptake by 27–49% relative to vehicle (*p* < 0.001; **Fig. 3a**; **Fig. S3**). Importantly, treatment effects were detectable by 3 dpi, when uptake was reduced by 38.0% with FPV (*p* = 0.008), 28.7% with RBV (*p* = 0.04), and 53.2% with BXV (*p* = 0.002; **Fig. 3b**). SUV measurements were less sensitive to these early differences, likely because of disease-associated weight loss (**Fig. S4**). *Ex vivo* gamma counting confirmed substantially reduced lung [^18^F]FDG retention following antiviral treatment (68.1–88.6%; *p* < 0.001; **Fig. S5**).

**Fig. 3.**
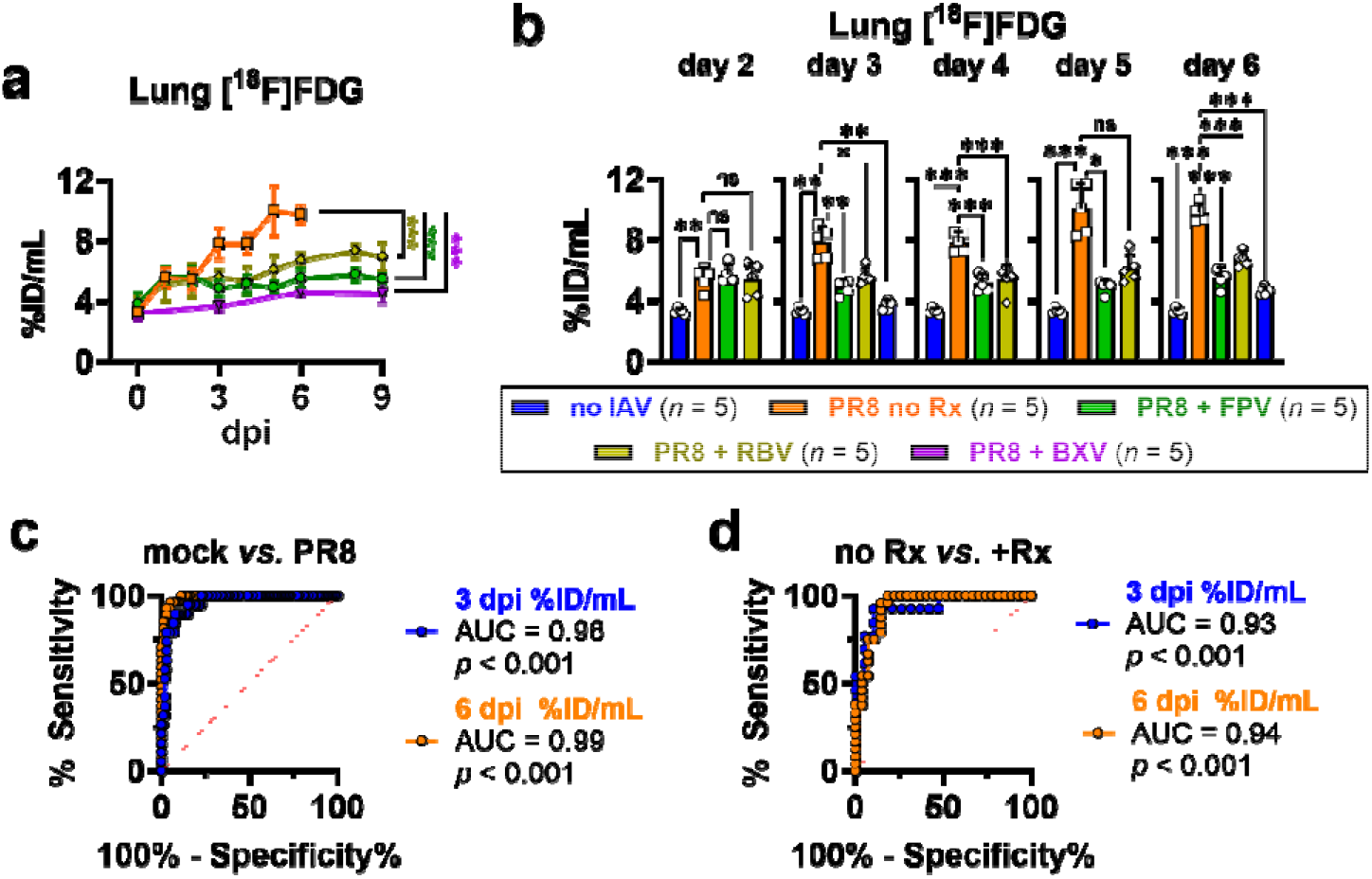
Evaluation of lung [^18^F]FDG uptake as a biomarker of influenza-associated lung inflammation and predictor of survival following antiviral therapy. a, Temporal profile of lung [^18^F]FDG uptake in mice infected with virus and treated with favipiravir (FPV), ribavirin (RBV), or baloxavir marboxil (BXV); or vehicle control (No Rx) over 9 days of infection. b, Comparison of lung uptake on every day from 2–6 days post-infection (dpi). c-d, Receiver-operating characteristic (ROC) curves of sensitivity and specificity of lung [^18^F]FDG uptake as h, biomarker of lung infection and inflammation; and as i, pharmacodynamic biomarker of antiviral treatment in influenza that predicts survival beyond 6 dpi. AUC (*c*-value) for lung uptake expressed as %injected tracer per tissue volume (%ID/mL) at 3 dpi and 6 dpi are compared. Data are presented as mean ± s.d. (*n* = 5 mice per group) representative of three independent experiments and compared using one-way ANOVA with Tukey’s correction. * *p* < 0.05; ** *p* < 0.005; ** *p* < 0.001; *ns*, not significant.

### Lung [^18^F]FDG uptake predicts influenza mortality and antiviral response

To evaluate the predictive performance of PET imaging biomarkers, receiving operator characteristic (ROC) analysis was conducted. This demonstrated that lung [^18^F]FDG uptake strongly predicted influenza lethality at both 3 dpi (AUC = 0.96) and 6 dpi (AUC = 0.99; *p* < 0.001; **Fig. 3c**). Thresholds of >4.68 %ID/mL at 3 dpi and >5.18 %ID/mL at 6 dpi exhibited high positive predictive values (PPV) of infection-associated lethality (PPV = 0.98 for 3 dpi and PPV = 0.99 for 6 dpi) with high sensitivity (100%) and specificity (71-85%; **Tables S1,S2**).

[^18^F]FDG uptake also predicted survival following antiviral treatment (AUC = 0.93 at 3 dpi and 0.94 at 6 dpi; *p* < 0.001; **Fig. 3d**). At 3 dpi, lung uptake <5.35 %ID/mL predicted survival with 92.3% sensitivity and 89.5% specificity (PPV = 0.97), while <7.53 %ID/mL uptake at 6 dpi achieved 100% sensitivity and 81.5% specificity (PPV = 0.96; **Tables S3,S4**). SUV measurements showed lower predictive accuracy (**Fig. S6**), supporting lung [^18^F]FDG uptake expressed as %ID/mL as the more sensitive pharmacodynamic and prognostic measure.

### Metabolic lung volume sensitively detects antiviral response and predicts disease outcome

To capture the spatial dynamics of influenza-associated inflammation, metabolic lung volumes (MLV) were measured. MLV, defined as lung volume exceeding a baseline [^18^F]FDG uptake threshold, progressively increased with disease in vehicle-treated mice but was reduced by 40.5–74.6% with antiviral therapy (*p* < 0.001; **Fig. 4a**; **Fig. S7**). As early as 3 dpi, MLV was reduced by 59.7% (*p* < 0.001), 33.8% (*p* = 0.006), and 75.0% (*p* < 0.001) with FPV, RBV, and BXV, respectively (**Fig. 4b**).

**Fig. 4.**
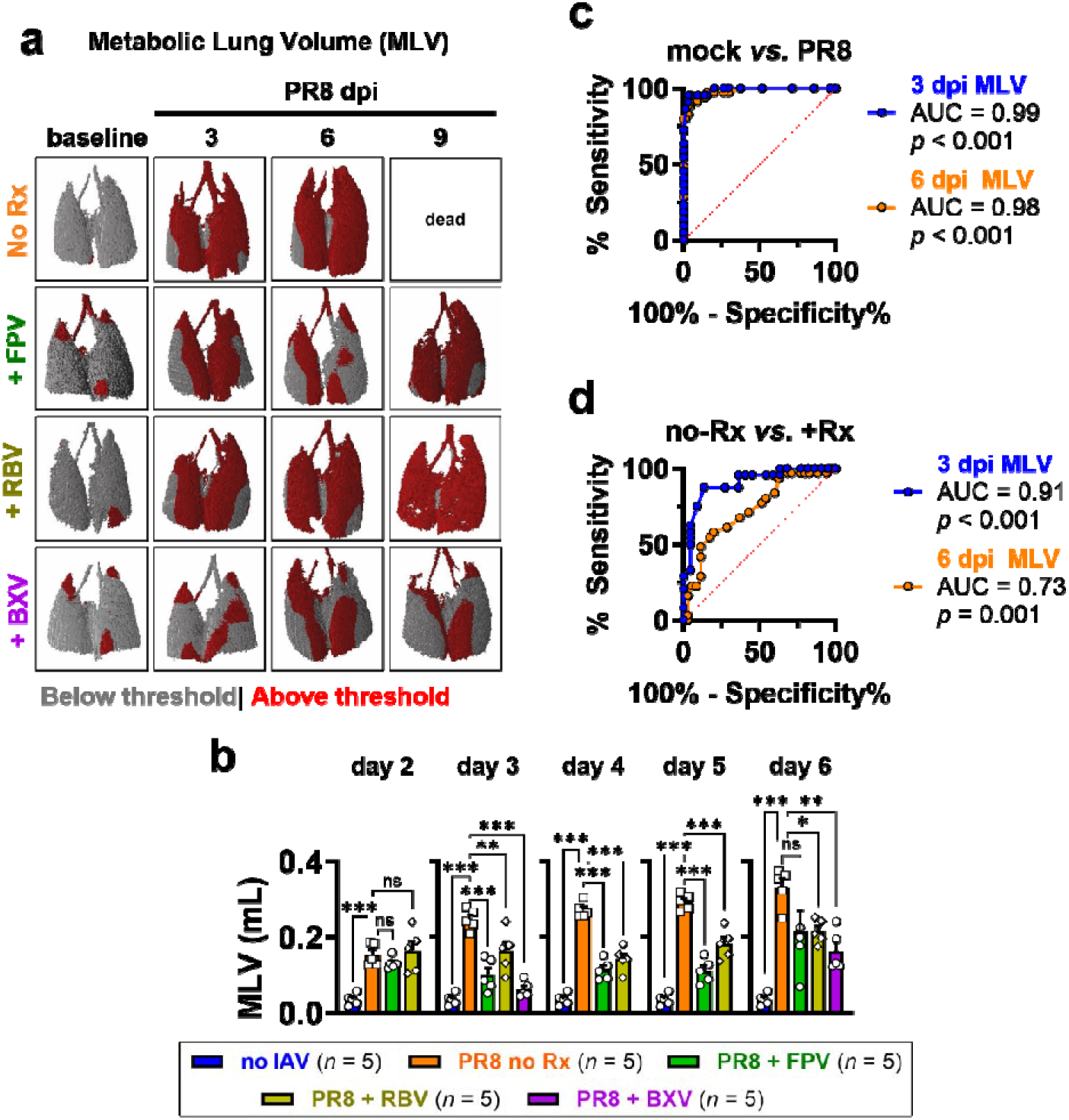
Evaluation of metabolic lung volume (MLV) as a biomarker of lung inflammation and predictor of survival following antiviral therapy. a, Representative 3D surface-rendered images of MLV depicting basal [^18^F]FDG uptake in grey and elevated uptake in red. b, Comparison of MLV in mice infected with virus and treated with favipiravir (FPV), ribavirin (RBV), or baloxavir marboxil (BXV); or vehicle control (No Rx) at 2-6 days post-infection (dpi). c-d, ROC curves of sensitivity and specificity of MLV as c, inflammation biomarker of infection-associated inflammation; and d, pharmacodynamic biomarker of antiviral treatment in influenza, predicting survival beyond 6 dpi. AUC (*c*-value) for MLV at 3 dpi and 6 dpi are compared. Data in *b* are presented as mean ± s.d. (*n* = 5 mice per group), representative of three independent experiments. Means were analyzed by one-way ANOVA with Tukey’s correction for comparison against vehicle control group. * *p* < 0.05; ** *p* < 0.005; ** *p* < 0.001; *ns*, not significant.

MLV accurately predicted influenza lethality at 3 and 6 dpi (AUC = 0.99 and 0.98; *p* < 0.001; **Fig. 4c**). MLV >0.075 mL at 3 dpi accurately estimated mortality with 100% sensitivity (PPV = 0.99) and 79.7% specificity (NPV = 1.0), while MLV >0.045 mL at 6 dpi similarly achieved highly reliable estimations (NPV = 1.0, PPV = 0.98; **Table S5-S6**). MLV also accurately predicted survival following antiviral therapy, particularly at 3 dpi (AUC = 0.91; *p* < 0.001), where MLV <0.185 mL estimated survival with 95.8% sensitivity (PPV = 0.91) and 63.6% specificity (NPV = 0.79; **Fig. 4d**; **Table S7**). MLV at 6 dpi provided comparably accurate survival estimates (AUC = 0.73, *p* = 0.001), particularly at MLV <0.255 mL (NPV = 0.71, PPV = 0.85; **Fig. 4d**; **Table S8**).

Thus, both lung [^18^F]FDG uptake and MLV provide early measures of antiviral pharmacodynamic response and disease outcome.

### Lung [^18^F]FDG-PET reflects suppression of pulmonary inflammation and viral burden

To evaluate the extent to which the imaging biomarkers reflect underlying lung inflammatory changes associated with antiviral treatment, inflammatory cytokine and chemokine expression at peak disease (6 dpi in No-Rx; 9 dpi for FPV, RBV, and BXV) were measured. At peak disease, antivirals broadly suppressed pulmonary inflammation, including neutrophil mediators (MCP-1, IL-7, MIP-1α, LIF) and systemic inflammatory cytokines (IL-6, IL-17E, IL-17A); while RBV resulted in more potent downregulation of several inflammatory mediators (IL-13, IL-7Rα, RANKL, IL-5, IL-33R; **Fig. 5a,b**; **Fig. S8,S9**). Lung [¹ F]FDG uptake strongly correlated with MLV (*r* = 0.93, *p* < 0.001) and with viral burden and inflammatory mediators including MCP-1, IL-6, MCP-3, IL-3, and MIP-1α (**Fig. 5c**; **Fig. S10**). MLV showed similar associations with viral load and inflammatory cytokines (**Fig. S11**), linking PET-derived measures to the underlying virologic and inflammatory response.

**Fig. 5.**
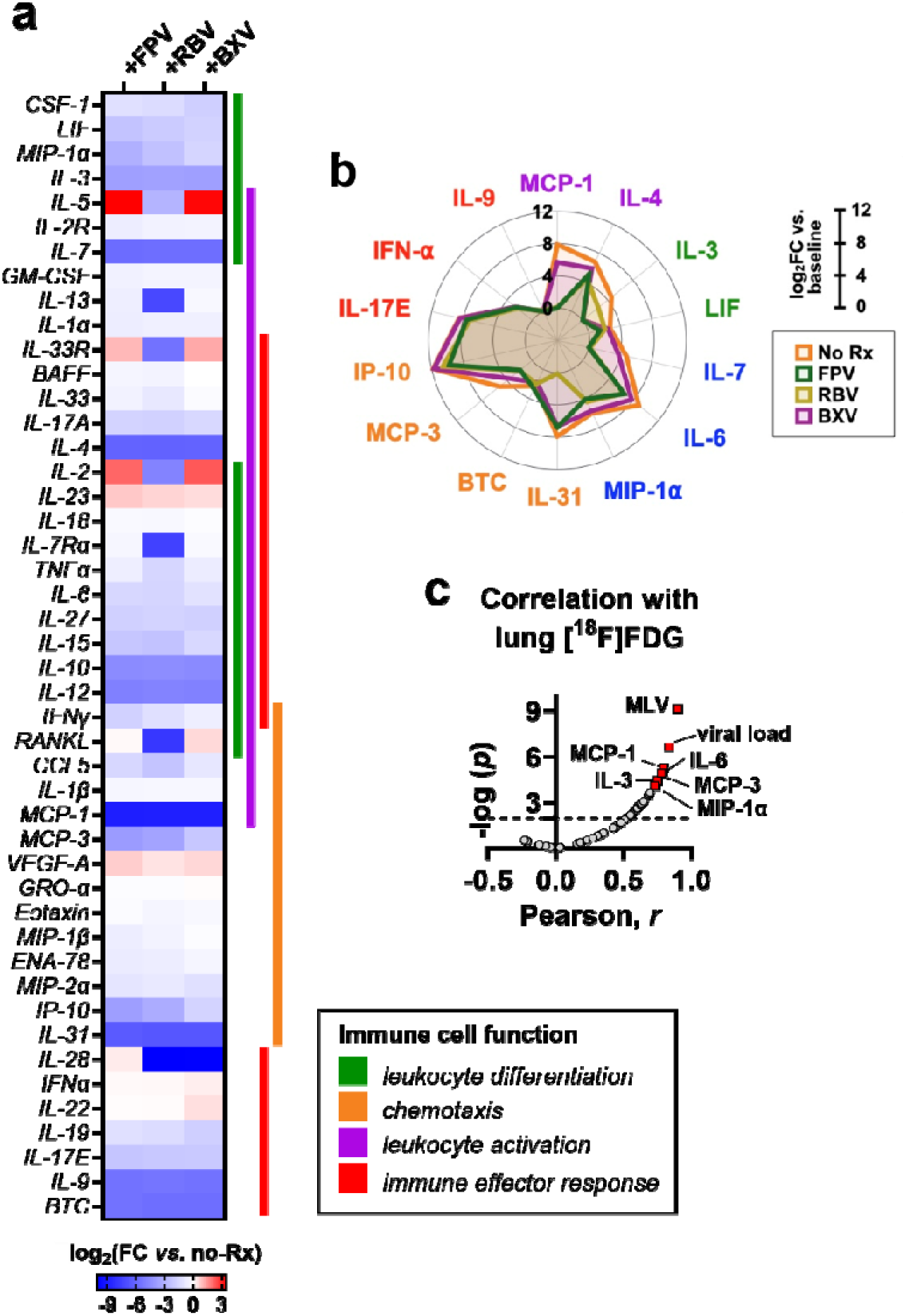
Association of lung [^18^F]FDG uptake with inflammation markers in PR8-infected mice treated with antivirals. **a,** Heatmap of selected cytokines and chemokines involved in immune cells differentiation, activation, chemotaxis and effector response. Mice were treated with antivirals favipiravir (FPV), ribavirin (RBV) or baloxavir marboxil (BXV); on vehicle control (no Rx). Data are shown as log_2_ fold-change expression relative to vehicle-treated mouse lungs (log_2_FC *vs.* no-Rx). Proteins associated with leukocyte activation (*purple*), differentiation (*green*), or both (*blue*) are indicated. Proteins involved in immune effector response are shown in *red*. **b,** Radar plots of the twelve most downregulated proteins in the lungs, shown as log_2_FC expression relative to healthy, uninfected lungs (baseline). **c,** Volcano plot depicting correlations between lung [^18^F]FDG uptake and infection and inflammation parameters. Horizontal dashed lines indicate the threshold of significance (*p* < 0.05).

### Validation in pandemic H1N1/CA-09 infection

To assess the translational relevance of lung [^18^F]FDG-PET as a pharmacodynamic biomarker we repeated the study in mice infected with the clinical isolate H1N1/CA-09, the representative strain of the 2009 H1N1 pandemic virus, and treated with BXV (**Fig. 6a**). BXV prolonged survival (OSm >9 *vs*. 5 days; *p* < 0.001), reduced weight loss, suppressed inflammatory cytokines (MCP-1, LIF, MIP-1α, IFN-α, IL-2, IL-6, TNF-α) and reduced lung viral replication by 30% relative to vehicle-treated lungs (8.7×10^4^ TCID_50_ *vs*. 1.7×10^7^ TCID_50_, *p* = 0.03; **Fig. 6b–d**; **Fig. S12,S13**).

**Fig. 6.**
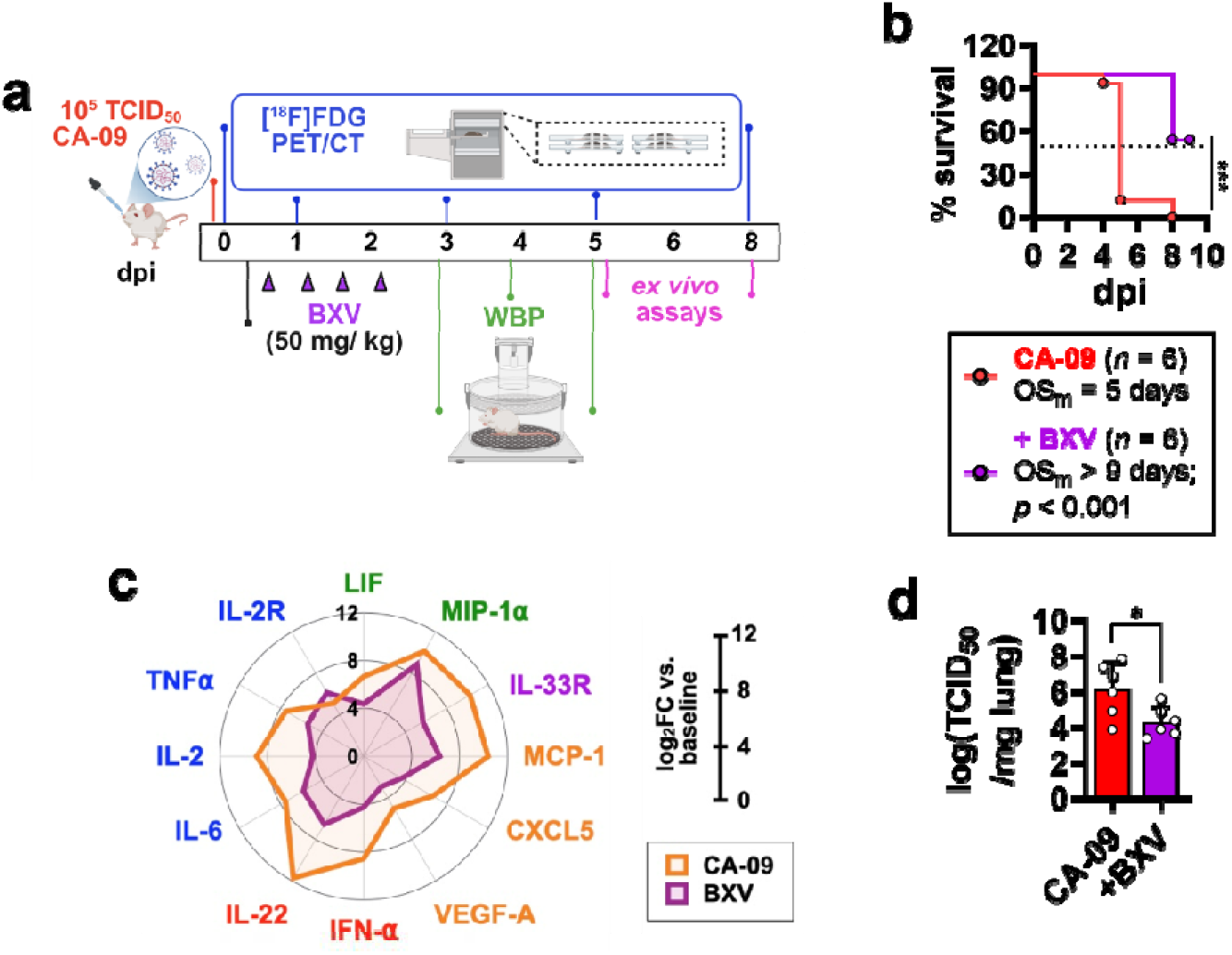
Lung [^18^F]FDG-PET is an early biomarker of baloxavir treatment in CA-09-infected mice. **a,** Study timeline showing virus inoculation, baloxavir marboxil (BXV) treatment, whole-body plethysmography (WBP), and experimental endpoints. CA-09 infection was performed on day 0, and BXV was administered twice daily by oral gavage (50 mg/kg per dose). Serial [^18^F]FDG-PET/CT imaging was performed at 0, 1, 3, and 5 days post-infection (dpi) in untreated mice and extended to 8 dpi in BXV-treated mice. WBP measurements were conducted at 3–5 dpi. *Ex vivo* analyses, including histopathology and protein ELISA, were performed at study endpoints. **b,** Kaplan–Meier survival curves. Dotted line indicates the humane endpoint (70% of initial body weight). Survival was compared using the log-rank (Mantel–Cox) test. **c,** Body weight changes following infection. **d,e,** Representative serial coronal slices of **d,** [^18^F]FDG-PET/CT and **e,** CT images. Lung volumes of interest (VOIs) are indicated by red dashed lines; *hrt*, heart. *lng*, lung. *bm*, bone marrow. **f–h,** Quantification of **f,** lung [^18^F]FDG uptake (%ID/mL), **g,** metabolic lung volume (MLV) from PET, and **h,** CT-derived healthy lung volume. **i,** Lung viral load measured at necropsy (5 dpi for untreated mice; 8 dpi for BXV-treated mice). Data are mean ± s.d. Statistical analysis was performed using Welch’s *t*-test. **j-k**, Receiver-operating characteristic (ROC) curves of sensitivity and specificity of lung [^18^F]FDG uptake as **j**, inflammation biomarker of infection; and **i**, pharmacodynamic biomarker of antiviral treatment in influenza. AUC for uptake at 3 dpi and 6 dpi are compared. * *p* < 0.05; ** *p* < 0.005; *** *p* < 0.001; ns, not significant.

Serial PET detected treatment response by 3 dpi. Compared with vehicle, BXV reduced lung [^18^F]FDG uptake by 37.4% (*p* = 0.02) and MLV by 67.5% (*p* < 0.001; **Fig. 7a,b**; **Fig. S15**). In contrast, CT-derived healthy lung volume did not distinguish treatment groups until 5 dpi (**Fig. 7b**).

**Fig. 7.**
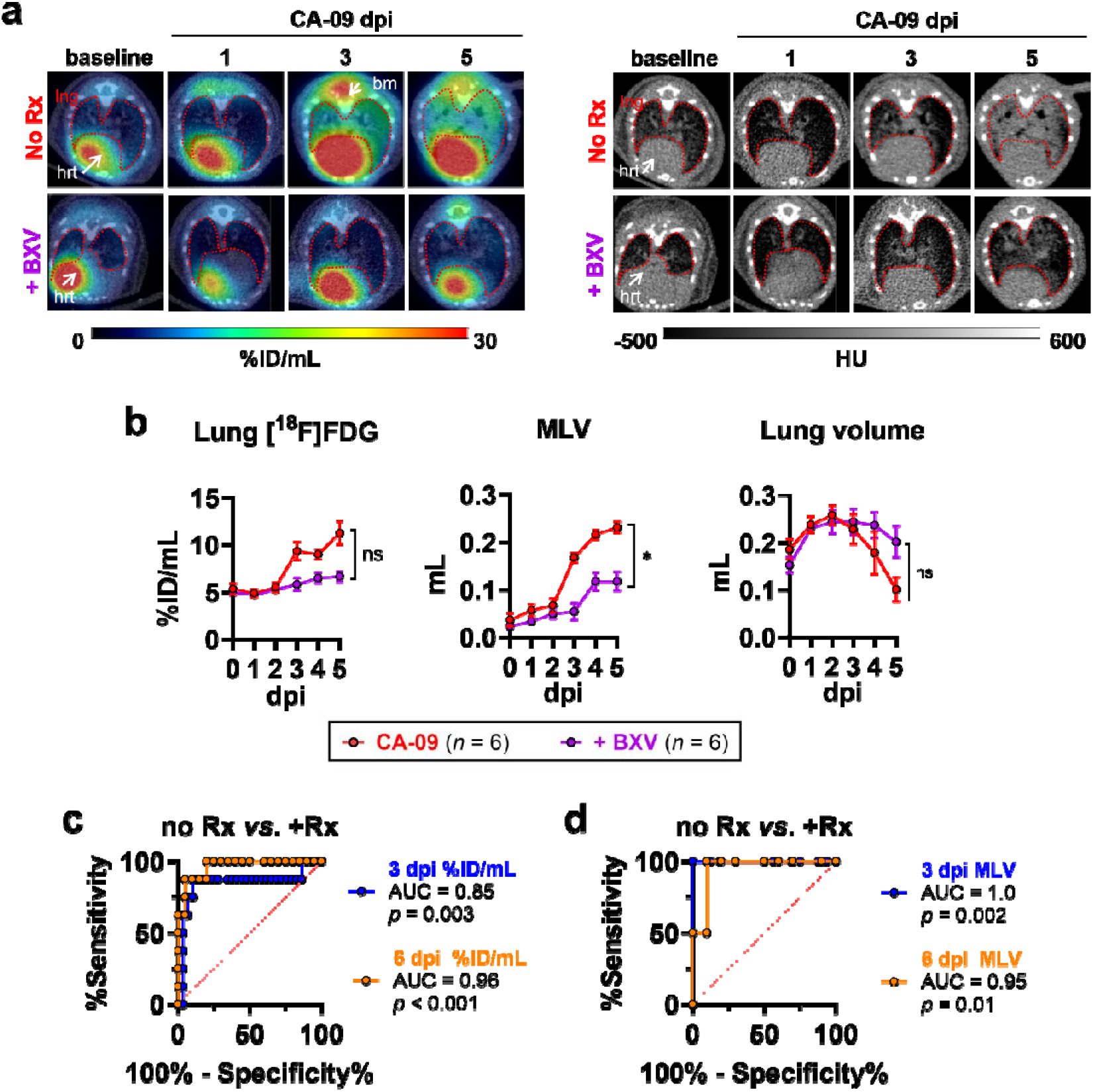
Validation of lung [^18^F]FDG uptake and metabolic lung volume (MLV) as inflammation biomarker and predictor of survival in CA-09-infected mice. a, Representative serial coronal slices of lung [^18^F]FDG-PET/CT and CT images. Lung volumes of interest (VOIs) are indicated by red dashed lines; *hrt*, heart. *lng*, lung. *bm*, bone marrow. b, Quantification of lung [^18^F]FDG uptake (%ID/mL), metabolic lung volume (MLV), and CT-derived healthy lung volume. Data are mean ± s.d from 6 mice per group representative of 3 independent studies. c-d, Receiver-operating characteristic (ROC) curves of sensitivity and specificity of lung c, [^18^F]FDG uptake; and d, MLV as pharmacodynamic biomarker of antiviral treatment in influenza. AUC for uptake at 3 dpi and 6 dpi are compared. Statistical analyses in *b* were using conducted Welch’s *t*-test. * *p* < 0.05; ** *p* < 0.005; *** *p* < 0.001; *ns*, not significant.

Lung [^18^F]FDG uptake predicted therapeutic response and survival at 3 dpi (AUC = 0.85, *p* = 0.003) and 5 dpi (AUC = 0.96, *p* < 0.001; **Fig. 7c**). Lung uptake <5.47 %ID/mL at 3 dpi predicted survival with 87.5% sensitivity (PPV = 0.97) and 89.7% specificity (NPV = 0.64); while <7.19 %ID/mL uptake at 6 dpi resulted in similarly accurate survival estimations (PPV = 0.95, NPV = 1.0; **Table S9,S10**). Lung [^18^F]FDG uptake also correlated strongly with viral load and inflammatory mediators including MCP-1, LIF, and MIP-1α (**Fig. S16**).

MLV similarly predicted treatment response and survival (AUC = 0.10, *p* = 0.002 at 3 dpi; AUC = 0.95, *p* = 0.01 at 5 dpi) and correlated with viral burden and inflammatory cytokines (**Fig. 7d**; **Fig. S17**). MLV <0.085 mL at 3 dpi predicted survival with 100% sensitivity and specificity (NPV = 1.0, PPV = 1.0), while MLV <0.14 mL at 5 dpi resulted in comparable estimates (PPV = 0.98, NPV = 1.0 **Table S11,S12**). These results validate [^18^F]FDG-PET-derived measures as pharmacodynamic and prognostic biomarkers across two H1N1 models.

### [^18^F]FDG-PET detects antiviral response earlier than respiratory function testing

To assess whether conventional lung function testing could distinguish the effects of antiviral treatment in this model, whole-body plethysmography (WBP) in awake mice was performed (**Fig. 6a**). WBP and AUC analysis throughout the 10-min data recording detected respiratory impairment in CA-09 infected mice *vs.* healthy mice—including prolonged inspiratory and expiratory pauses (EIP and EEP) and altered breathing-transition parameters (Penh, PAU, T_P_), reflective of bronchoconstriction (**Fig. 8a**; **Fig. S18,S19**). Penh was 2.2-fold (*p* < 0.001) and 2.0-fold (*p* = 0.001) higher than control lungs at 3 dpi and 4 dpi, respectively; while PAU was elevated 2.0-fold (*p* < 0.001) and 1.6-fold (*p* < 0.001) (**Fig. 8a**).

**Fig. 8.**
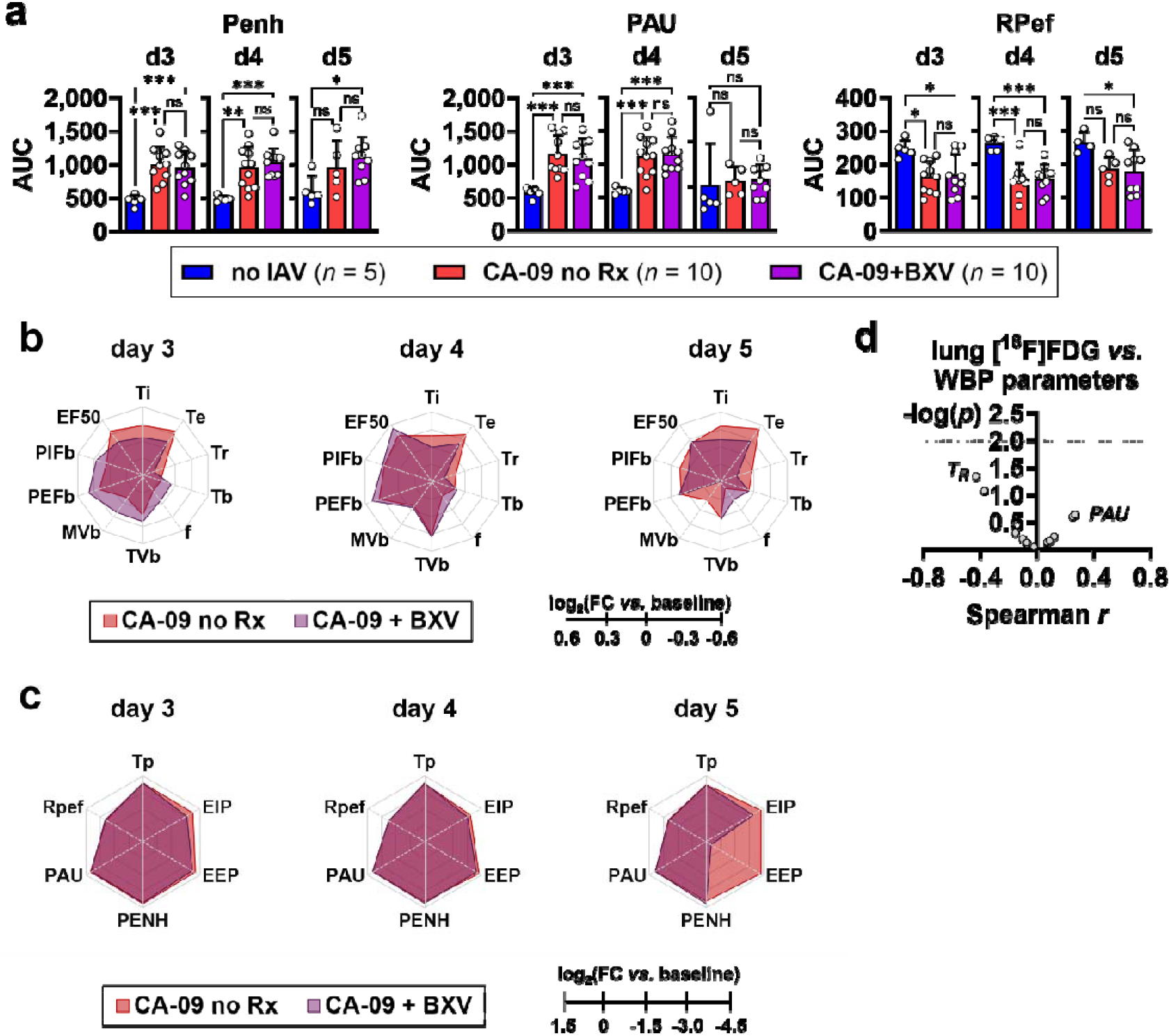
Evaluation of respiratory parameters by whole-body plethysmography (WBP) following antiviral treatment. **a,** Comparison of WBP parameters enhanced pause (Penh); pause (PAU); and RPef between mice infected with A/CA-09 and treated with baloxavir marboxil (BXV) or vehicle (no Rx). Data are shown as mean ± s.d. of AUC over 10-min data acquisition period. **b,c**, Radial plot of WBP parameters depicting log_2_ fold-change AUC relative to healthy lungs (log_2_FC *vs*. baseline) from days 3 to 5 post-infection. **d**, Volcano plot depicting correlations between lung [^18^F]FDG uptake and WBP parameters in virus-infected and mock-infected lungs from days 3-5 post-infection. Horizontal dashed lines indicate the threshold of significance (*p* < 0.05). *r*, Spearman coefficient. Means in ***a*** were compared with one-way ANOVA with Tukey’s correction. * *p* < 0.05; ** *p* < 0.005; *** *p* < 0.001; *ns*, not significant. Details of WBP parameters could be found in Fig. S18.

However, although these measures distinguished virus-infected from healthy mice, significant differences between BXV- and vehicle-treated animals were largely restricted to 5 dpi (**Fig. 8b,c**). Here, BXV reduced EIP and EEP by 43.3% (*p* < 0.001) and 96.7% (*p* = 0.005), respectively (**Fig. S19**). Other WBP parameters did not effectively distinguish treatment groups, and WBP measurements did not correlate with lung [^18^F]FDG uptake (**Fig. 8d**; **Fig. S19**).

Together, these findings demonstrate that lung [^18^F]FDG uptake and MLV detect antiviral efficacy by 3 dpi, preceding detectable structural changes by CT and treatment-associated improvements in respiratory function.

## Discussion

In this study, we identify lung [^18^F]FDG-PET as a sensitive pharmacodynamic biomarker of antiviral efficacy in influenza A virus infection. Across two murine H1N1 models and three mechanistically distinct antivirals, serial PET detected suppression of pulmonary inflammation as early as 3 days post infection, preceding CT-detectable structural abnormalities and changes in respiratory function. PET measures of lung [^18^F]FDG uptake and metabolic lung volume (MLV) correlated with pulmonary cytokine responses and tissue viral burden, linking the imaging signal to both host inflammation and viral replication. By contrast, whole-body plethysmography was less sensitive to early treatment effects. These findings indicate that [^18^F]FDG-PET can detect early treatment-responsive changes in pulmonary metabolism before structural or physiological abnormalities become apparent.

Our findings extend evidence supporting molecular imaging for monitoring host-pathogen interactions and therapeutic response [23, 24, 34]. As demonstrated in tuberculosis, PET enables longitudinal tissue-level assessment that is difficult to achieve with conventional sampling [25]. Influenza may similarly benefit from an imaging biomarker that captures treatment effects directly within the lung. The established clinical use of [^18^F]FDG-PET also provides a potential pathway for translation from preclinical models to clinical antiviral studies.

Among the PET metrics evaluated, MLV was the most sensitive to antiviral treatment, potentially because it captures the spatial extent of metabolically active lung tissue. Its ability to distinguish treated from untreated animals before CT-detectable structural changes suggests that metabolic imaging may provide an earlier indicator of therapeutic response than conventional measures of lung injury. This distinction may be particularly relevant when evaluating interventions intended to suppress viral replication before irreversible pulmonary injury develops.

Although all three antivirals reduced pulmonary inflammation and viral burden, cytokine profiling revealed treatment-specific host responses. Favipiravir and baloxavir produced broadly similar signatures consistent with reduced viral replication and downstream innate immune activation, whereas ribavirin increased ST2, IL-5, IL-2, and RANKL despite broader suppression of pro-inflammatory mediators. These findings are consistent with the reported immunomodulatory effects of ribavirin [35–37] and demonstrate that similar reductions in PET signal can accompany distinct immunological responses. Integrating PET with immune profiling may therefore help distinguish antiviral effects on pulmonary inflammation from specific changes in host immunity.

Baloxavir produced the earliest and largest reductions in pulmonary [^18^F]FDG signal in our models. Clinically, baloxavir has been associated with reduced influenza transmission, consistent with rapid suppression of viral replication and shedding [15, 38]. Together, these findings support evaluation of PET-derived measures alongside virological and clinical endpoints in early-phase antiviral studies, particularly when viral and host responses may not be closely coupled [39].

Importantly, however, whether PET responses in experimental influenza predict therapeutic efficacy or clinical outcomes in humans remains unknown.

Several limitations should be considered. Murine models do not fully reproduce the heterogeneity of human influenza, and the timing and magnitude of PET responses require validation in other relevant models and ultimately in humans. [^18^F]FDG is also a non-specific marker of glucose metabolism and cannot distinguish the contributions of recruited immune cells, resident pulmonary cells, and virus-induced metabolic changes. More cell-selective tracers (*e.g.*, myeloid-, T cell-, or TSPO-directed probes) and kinetic imaging may improve biological specificity [40]. Finally, experiments were performed exclusively in female BALB/c mice; studies across sex, age, and clinically relevant comorbidities (*i.e.,* asthma, COPD, diabetes, pregnancy) will be required to establish generalizability.

In conclusion, lung [¹ F]FDG-PET provides an early, quantitative measure of antiviral response in experimental influenza A infection that reflects pulmonary inflammation and viral burden before structural or physiological changes become apparent. These findings establish treatment responsiveness of the imaging signal and provide a rationale for evaluating [^18^F]FDG-PET as a pharmacodynamic endpoint in translational influenza studies.

## Methods

### Ethics statement

All animal experiments were conducted in accordance with protocols approved by the Institutional Animal Care and Use Committee (IACUC) of Duke-NUS Medical School and SingHealth (approval no. 2020/SHS/1607).

### Materials

Female BALB/c mice (3–4 weeks old) were obtained from In Vivos (Singapore) and housed in a biosafety level 2 (BSL-2) facility under specific pathogen–free conditions.

Influenza A/Puerto Rico/8/34 (H1N1; PR8) was obtained from the American Type Culture Collection (ATCC VR-1469). Influenza A/California/04/09 (H1N1; CA-09) was generated by reverse genetics and provided by Prof. Gavin Smith (Duke-NUS Medical School).

Antivirals with mechanistically distinct modes of action were evaluated in this study: Favipiravir (FPV), an RNA-dependent RNA polymerase inhibitor; Baloxavir (BXV), a cap-dependent endonuclease inhibitor; and Ribavirin (RBV), a broad-spectrum nucleoside analogue. FPV (HY-14768), BXV (HY-109025A), and RBV (HY-B0434) were purchased from MedChemExpress (USA) at >99.97% purity.

The radiotracer [^18^F]FDG (>95% radiochemical purity) was obtained from Singapore Radiopharmaceuticals (Singapore).

### Study design

Two murine models of influenza A/H1N1 infection models were used in this study: one using the mouse-adapted PR8 strain and another using the clinical isolate CA-09. Mice infected with either virus were given twice-daily treatment with either antivirals or vehicle control commencing at 6 h post-infection. Mice were subjected to daily serial [^18^F]FDG-PET/CT imaging daily from pre-infection to day 9 post-infection. Tissues were collected for *ex vivo* assays at study endpoint (day 6 for mice treated with vehicle control and day 9 for antiviral-treated animals) and paired with the final day of PET scan.

### Virus infections

Mouse infection with influenza was conducted as described [33]. Animals were anesthetized by intraperitoneal (i.p.) injection with ketamine (80 mg/kg) and xylazine (15 mg/kg) cocktail and subsequently instilled with 30 µL mouse-adapted PR8 virus (1×10^6^ TCID_50_) or clinical isolate CA-09 virus (1×10^5^ TCID_50_) into the left nostril. Mice were kept in dorsal recumbence throughout the procedure and until the animals awakened.

### Antiviral Treatment

Drugs were reconstituted in absolute dimethyl sulfoxide (DMSO) and subsequently formulated for oral administration following the manufacturer’s protocol: 50 mg/kg in a suspension of 10% DMSO, 40% polyethylene glycol 300 (PEG 300), 5% Tween-80, and 45% saline. Drugs were administered by oral gavage twice daily (BID) within an 8h/16h dosing cycle, initiated at 6 h post-infection to model early therapeutic intervention and analogous to same-day antiviral initiation in high-risk patients presenting at the onset of influenza symptoms. This window was chosen based on evidence that the greatest clinical benefit of polymerase inhibitors is observed when treatment is initiated within the first 24 h of symptom onset [13, 14, 41], and is consistent with early treatment paradigms used in preclinical antiviral efficacy studies [42]. Treatment continued until day 4 post-infection for FPV, or day 2 post-infection for BXV and the broad-spectrum antiviral comparator RBV, following the study timeline shown in **Fig. 1a** and **Fig. 6a**.

### Positron emission tomography-computed tomography (PET-CT) imaging and *ex vivo* assays

PET/CT imaging studies were conducted as described previously [33]. After the final PET/CT scan (on either day 6 or 9 post-infection), mice were quickly euthanized by anaesthetic overdose (10 mg ketamine, 1 mg xylazine in 100 µL). Blood was removed by cardiac puncture, and the lungs were isolated. Full details can be found in Supplemental Files.

### Whole body plethysmography (WBP)

Pulmonary function was assessed non-invasively in conscious, non-anaesthetized, freely moving mice using whole-body plethysmography (WBP; Buxco DSI Finepointe, USA), which permits continuous respiratory monitoring without physical restraint or surgical instrumentation. Full details can be found in Supplemental Files.

### Statistical methods

All statistical analyses and data visualizations were performed using Prism version 10.2 (GraphPad Software, USA). Normality of data distribution was evaluated with Shapiro-Wilk test and verified in Q-Q plots. Equality of variance between groups was compared using Bartlett’s test. Data following Normal distribution and with equal variances from 2 groups or >2 groups were compared with Student’s *t*-test or one-way ANOVA, respectively; data with unequal variances were compared with either Welch’s *t-*test or Welch’s ANOVA. Multiple comparisons between groups in ANOVA were corrected using Tukey’s for equal variances, or Dunnett’s test for unequal variances.

Receiver operator characteristic (ROC) analyses to evaluate the predictive performance of imaging biomarkers were conducted in Prism. Positive predictive values (PPV) and negative predictive values (NPV) were calculated using theoretical fatality rates and treatment rates in the mouse model (high estimate of 95% and low estimate of 50%).

## Acknowledgement

The authors would like to thank Dr. Andrew Teo (Lee Kong Chian School of Medicine, Nanyang Technological University, Singapore) for performing Luminex assays; and Prof. Gavin Smith for providing the influenza A/H1N1/CA-09 strain generated by reverse-genetic engineering. We would also like to thank Professor Paul Tambayah (National University Hospital Singapore) for helpful discussions. This study was funded by the Singapore National Medical Research Council (NMRC) Open-Fund Large Collaborative Grant to the Integrated Innovations in Infectious Diseases (I3D) Programme, NMRC/OFLCG/MOH-000505–02 (to A-MC).

## Author contributions

CBLV and A-MC designed the study; AG performed PET/CT image acquisitions, while CBLV conducted all image analyses; CBLV, AG, SG, and JO performed *ex vivo* tissue biodistribution studies and assays for infection and inflammation. CBLV, JO and A-MC interpreted the results. CBLV and A-MC wrote the initial manuscript, and all authors contributed to revisions and approved the final version.

## Competing interests

The authors declare no competing interest regarding this study.

## Notes

### Competing Interest Statement

The authors have declared no competing interest.

