## Supplemental Information for "Lung [^18^F]FDG-PET is an early pharmacodynamic biomarker of antiviral efficacy in preclinical influenza"

**Extended Methods**

**Positron emission tomography (PET) and computed tomography (CT) imaging.** PET/CT imaging studies were conducted as described previously [1]. Mice anaesthetized with 2% isoflurane were intravenously injected with 30 MBq [^18^F]FDG and subjected to lung-focused static PET imaging for 20 min, followed by accurate whole-body CT scan for 2 min (tube current: 0.24 mA; tube voltage: 50 kV) (Vector^4^CT, MILabs, Japan). Four mice were simultaneously imaged. PET images were reconstructed at 0.8-mm voxel resolution using a SROSEM algorithm (16 subsets, 24 iterations, 1.5-μm Gaussian filter). PET and CT images were automatically co-registered, and PET attenuation correction was manually performed using the corresponding CT images.

Quantitative image analysis and image data visualization were carried out using PMOD software (v4.4, PMOD Technologies, Switzerland) as previously reported[1]. Volumes of interest (VOIs) were drawn on CT images using a semi-automated thresholding approach to segment lung space from surrounding tissues and refined using the cold iso-contouring function (HU ≤ -300 HU).

A second-degree analysis of lung metabolism by quantifying metabolic lung volume (MLV) as previously described [1]. MLV is defined as the tissue volume with [^18^F]FDG uptake higher than the baseline threshold, which is calculated as the mean $+$ (2$\times$σ) of lung uptake at pre-infection. Using PMOD, the MLV was calculated on PET images using the hot iso-contouring function with the calculated threshold as the minimum value.

***Ex vivo* assays.** After the final PET/CT scan (on either day 6 or 9 post-infection), mice were quickly euthanized by anaesthetic overdose (10 mg ketamine, 1 mg xylazine in 100 µL). Blood was removed by cardiac puncture, and the lungs were isolated. [^18^F]FDG retained in the tissues was measured by gamma counting and normalized to tissue weights. The lung left lobe was excised and fixed in 10% neutral buffered formalin for immunohistopathological examination, while the right lobes were flash-frozen in dry ice for RNA and protein quantification.

Immunohistopathological examination of the lungs was performed as previously described [2]. Freshly harvested lungs were fixed in 10% neutral buffered formalin, processed for paraffin embedding, and sectioned into 5 µm slices. Contiguous tissue sections were stained with hematoxylin & eosin (H&E) and subjected to immunofluorescence (I.F.) staining for viral nucleoprotein (Invitrogen, PA17221) and CD45 (Abcam, Ab10558). I.F. tissue sections were counterstained with DAPI, and autofluorescence was quenched using the with Vector^®^ TrueVIEW^®^ kit (Vector Laboratories, USA). Slides were imaged on a Nikon Ni-E inverted fluorescence microscope.

Flash-frozen lung tissues (~30 mg) were homogenized in phosphate-buffered saline (PBS, pH 7.4) containing a protease/phosphatase inhibitor cocktail (Sigma-Aldrich, USA). After centrifugation at >16,000 ⨉ g to clear the homogenate, the supernatant was supplemented with Triton X-100 to a final concentration of 1% (^v^/_v_) and subjected to one freeze-thaw cycle. Samples were clarified, and total protein concentrations were determined using a BCA assay. Cytokines and chemokines were measured using the Immune Monitoring 48-plex mouse ProcartaPlex panel (Thermo-Fisher, EPX480-20834-901) on a Luminex xMAP platform.

Total RNA was extracted from frozen lungs using RNeasy Mini Kit (Qiagen, Germany) according to the manufacturer’s instructions. Quantification of influenza A virus RNA using primers INFA-1 and INFA-2 as described previously [3] was performed by one-step reverse transcription quantitative PCR (RT-qPCR) Luna Universal One-Step RT-qPCR Kit (NEB, Cat. #E3005E). Absolute quantification of copy numbers was performed using influenza A/PR/8/34 genomic RNA standards (ATCC VR-1469DQ).

Determination of viable virus load in lung tissues was conducted using ELISA-based TCID_50_ assay as described by the WHO guidelines.

**Whole body plethysmography (WBP).** Pulmonary function was assessed non-invasively in conscious, non-anaesthetized, freely moving mice using whole-body plethysmography (WBP; Buxco DSI Finepointe, USA), which permits continuous respiratory monitoring without physical restraint or surgical instrumentation. Prior to measurements, female BALB/c mice underwent a 3-day acclimatization protocol in which each animal was placed individually in the plethysmography chamber for 30 min per day at a fixed time of day to minimise circadian variation and reduce stress-associated confounders[4]. On each experimental day, mice were acclimatized within the chamber for 20 min, followed by simultaneous recording of 17 respiratory parameters every 2 seconds, and continuously for 10 min in the unrestrained, conscious state.

Four parameters were extracted from the 10-min recording window: breathing frequency (f; breaths per minute, bpm), tidal volume (TVb; mL; reflecting the volume of air displaced per breath), pause (PAU, the ratio of expiratory to inspiratory time, reporting expiratory flow limitation), and enhanced pause (Penh, a dimensionless index reflecting the pressure waveform shape and used as a surrogate for airway resistance and bronchoconstriction). Using the raw values for each parameter, the area under the curve (AUC) was calculated across the recording period and used as the primary comparative metric between experimental groups, providing an integrated measure of respiratory function across the full recording window rather than reliance on single time-point values. All measurements were performed at the same time of day throughout the experimental period to minimise the effect 24-hour circadian rhythm differences on respiratory measurements.

**Supplementary Figures**

**Figure S1**

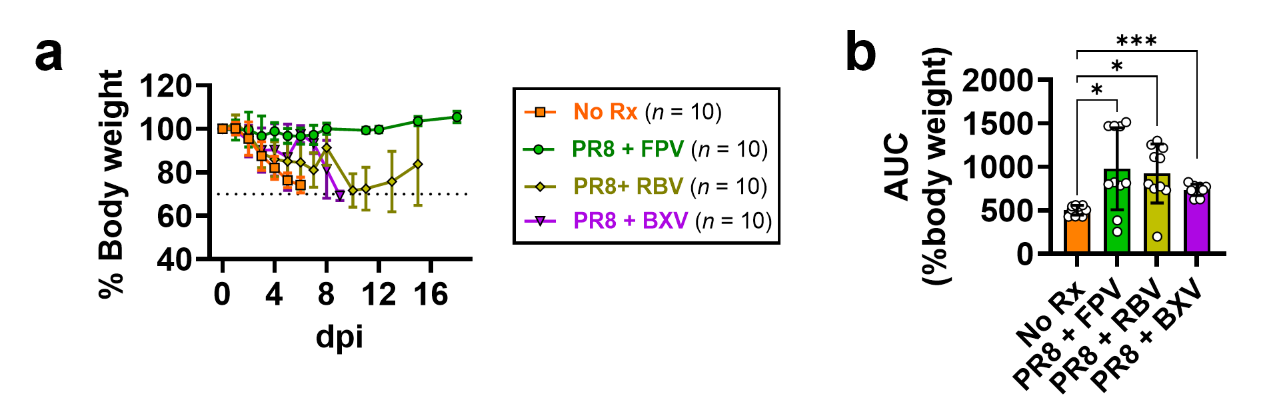

**Fig. S1 | Body weight changes following antiviral therapy in a murine model of influenza A/PR8 infection. a,** Body weights over 20 days reported as % of initial weight prior to virus infection. Dotted line indicates the humane endpoint (70% of initial body weight). **b**, Area under the curve (AUC) analysis of body weight changes. Data are shown as mean ± s.d. from 10 mice per group and compared using Welch’s ANOVA with Dunnett’s correction. * *p* < 0.05; ** *p* < 0.005; ** *p* < 0.001; *ns*, not significant.

**Figure S2**

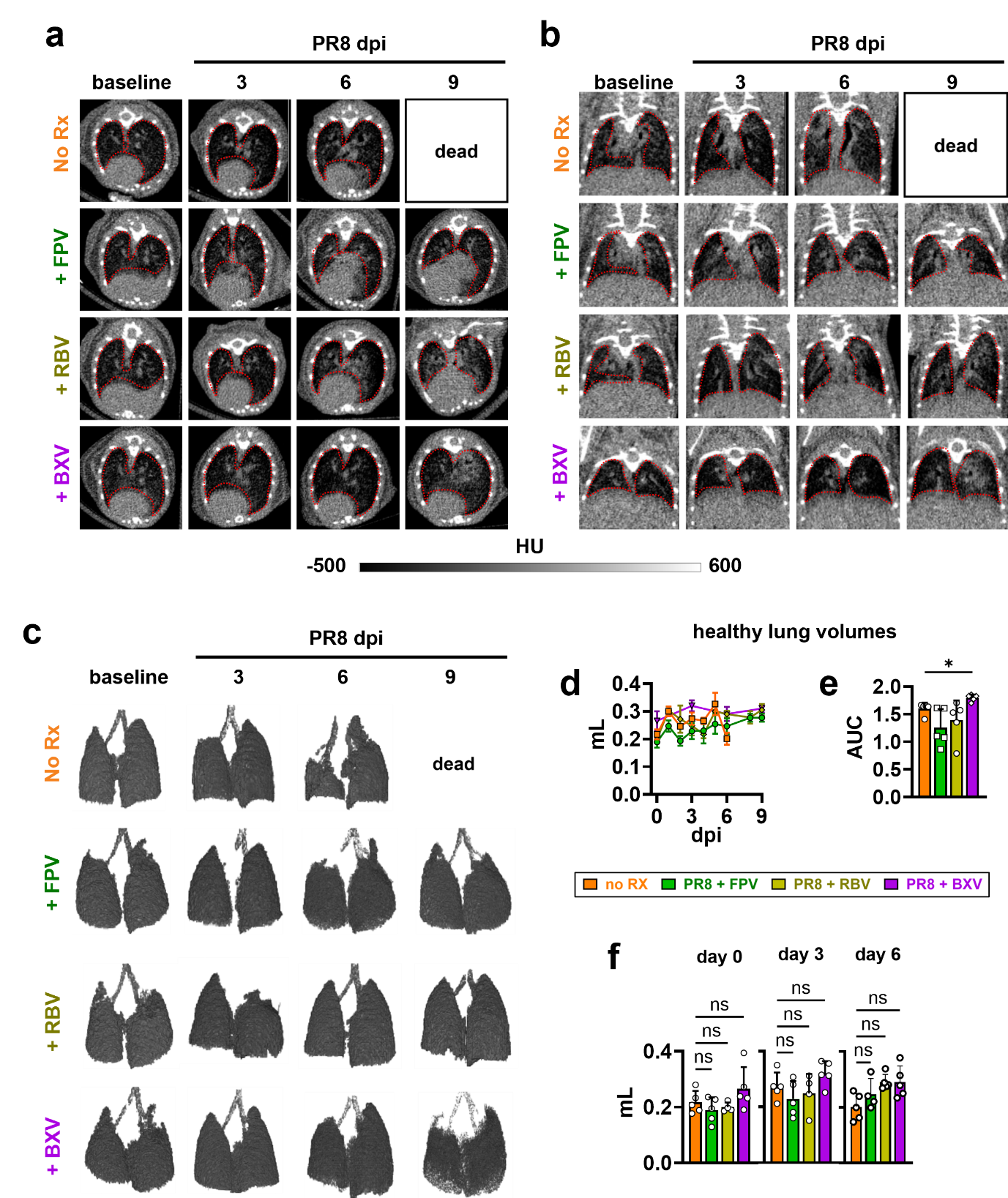

**Fig. S2 | Computed tomography (CT) imaging of lungs in influenza-infected mice. a,b,** Representative CT image slices in coronal (**a**) and transverse (**b**) views of mice infected with influenza A/H1N1/PR8 and either untreated (PR8 only) or treated with favipiravir (FPV), ribavirin (RBV), or baloxavir marboxil (BXV). **c,** Representative three-dimensional rendering of CT-derived healthy lung volumes. **d,** Temporal quantification of healthy lung volumes. **e,** Area under the curve (AUC) derived from **d**. **f,** Comparison of CT-derived lung volumes between groups at 0, 3, and 6 days post-infection (dpi). Data are from one experiment (*n* = 5 mice per group) and are representative of three independent studies. Values are mean ± s.d. Statistical analysis was performed using Welch’s ANOVA with Dunnett’s correction; * *p* < 0.05; ** *p* < 0.005; ** *p* < 0.001; *ns*, not significant.

**Figure S3**

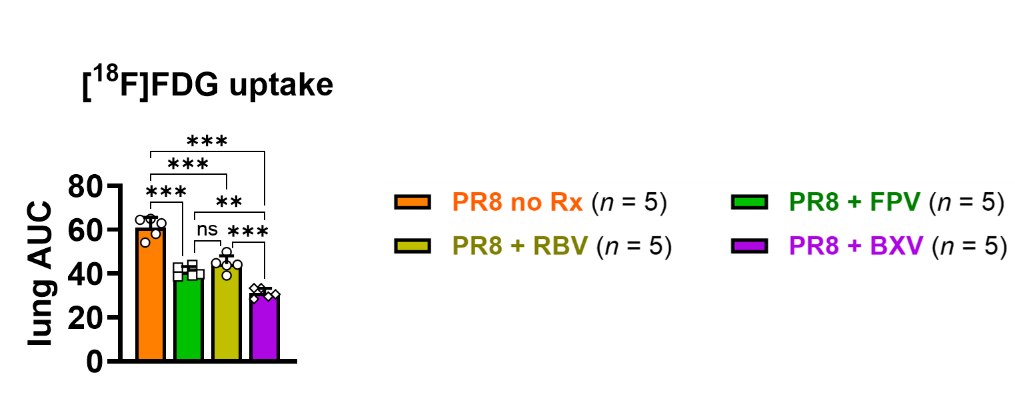

**Fig. S3 | AUC analysis of lung [^18^F]FDG uptake over 6 days of infection.** AUC of lung uptake calculated from days 0 to 6 post-infection. Mice were infected with virus and treated twice daily with favipiravir (FPV), ribavirin (RBV), or baloxavir marboxil (BXV); or vehicle control (No Rx). Data are shown as mean ± s.d. of 5 mice per group, representative of three independent experiments. Means were compared using one-way ANOVA with Tukey’s correction. * *p* < 0.05; ** *p* < 0.005; ** *p* < 0.001; *ns*, not significant.

**Figure S4**

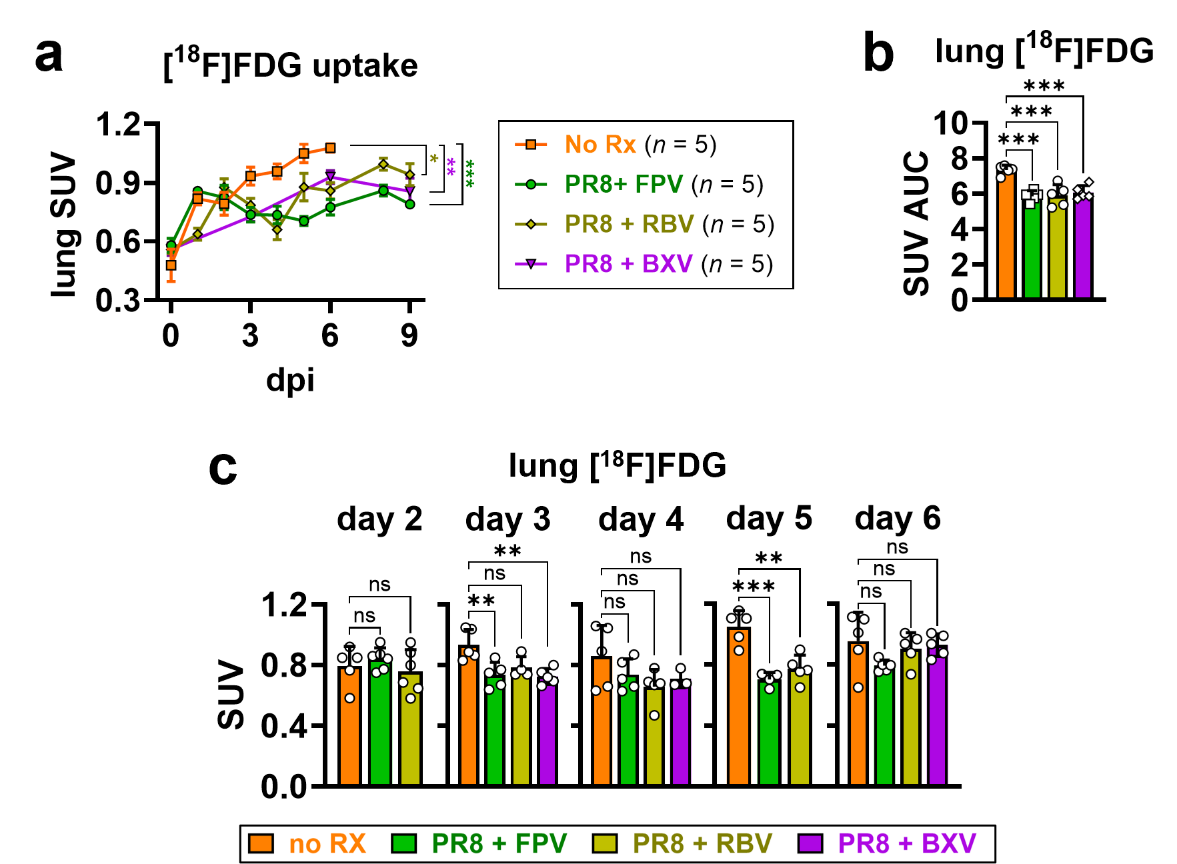

**Fig. S4 | Lung [¹⁸F]FDG uptake in PR8-infected mice quantified as standardized uptake values (SUV).** Mice were infected with virus and treated twice daily with favipiravir (FPV), ribavirin (RBV), or baloxavir marboxil (BXV); or vehicle control (No Rx) **a,** Temporal profile of lung [^18^F]FDG uptake expressed as SUV derived from serial PET imaging. **b,** Area under the curve (AUC) analysis calculated from ***a***. **c,** Comparison of lung SUV between groups at 2, 3, 4, 5, and 6 days post-infection (dpi). Data are mean ± s.d. from one experiment (*n* = 5 mice per group) and are representative of three independent studies. Statistical analysis was performed using one-way ANOVA with Tukey’s correction. * *p* < 0.05; ** *p* < 0.005; *** *p* < 0.001; *ns*, not significant.

**Figure S5**

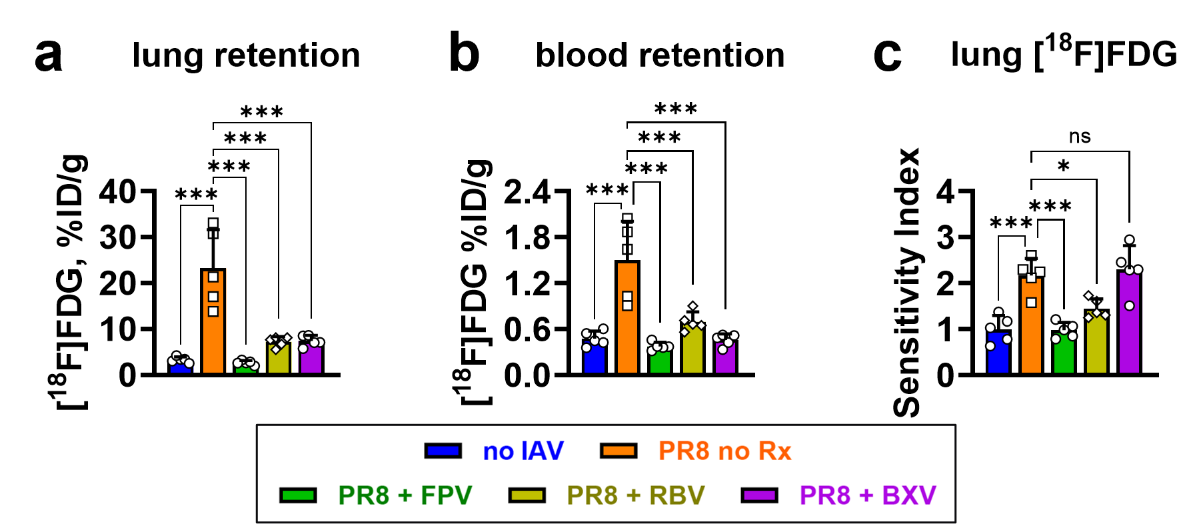

**Fig. S5 | *Ex vivo* validation of lung [¹⁸F]FDG uptake. a,b,** Gamma counting of **a,** lungs and **b,** blood harvested from mice euthanized immediately following the final PET scan. Mice were infected with virus and treated twice daily with favipiravir (FPV), ribavirin (RBV), or baloxavir marboxil (BXV); or vehicle control (No Rx). Mice without influenza infection (no IAV) are also included. Data are expressed as percentage injected dose per gram of tissue (%ID/g). **c,** Lung localization ratio, calculated as the lung-to-blood activity ratio in infected mice normalized to the corresponding ratio in mock-infected controls (no IAV). Data are from one experiment (*n* = 5 mice per group) and are representative of three independent studies. Values are mean ± s.d. Statistical analysis was performed using Welch’s ANOVA; * *p* < 0.05; ** *p* < 0.005; *** *p* < 0.001; *ns*, not significant.

**Figure S6**

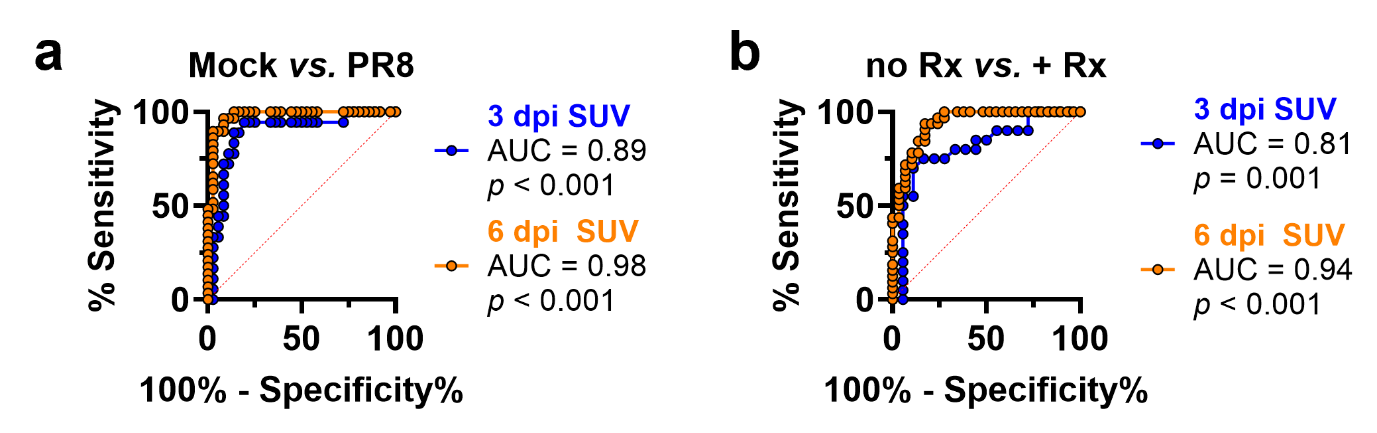

**Fig. S6 |** **Receiver-operating characteristic (ROC) analysis of lung [¹⁸F]FDG uptake quantified as standardized uptake values (SUV).** Evaluation of lung [^18^F]FDG SUV as **a**, biomarker of lung infection and inflammation; and as **b**, pharmacodynamic biomarker of antiviral treatment in influenza that predicts survival beyond 6 dpi.

**Figure S7**

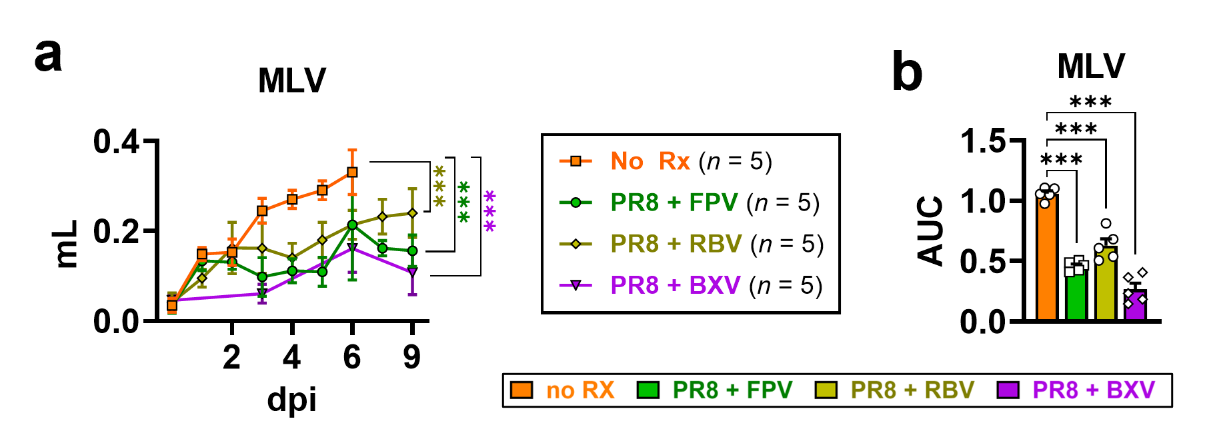

**Fig. S7 | Changes in metabolic lung volumes (MLV) following antiviral treatment in influenza A/PR8 model.** **a,** Quantification of MLV over time in mice infected with virus and treated with favipiravir (FPV), ribavirin (RBV), or baloxavir marboxil (BXV); or vehicle control (No Rx). **b,** AUC analysis of MLV in ***a***. Data are presented as mean ± s.d of *n* = 5 per group, representative of three independent experiments. Statistical analysis compared to vehicle treatment was performed using one-way ANOVA with Dunnett’s correction; * *p* < 0.05; ** *p* < 0.005; *** *p* < 0.001; *ns*, not significant.

**Figure S8**

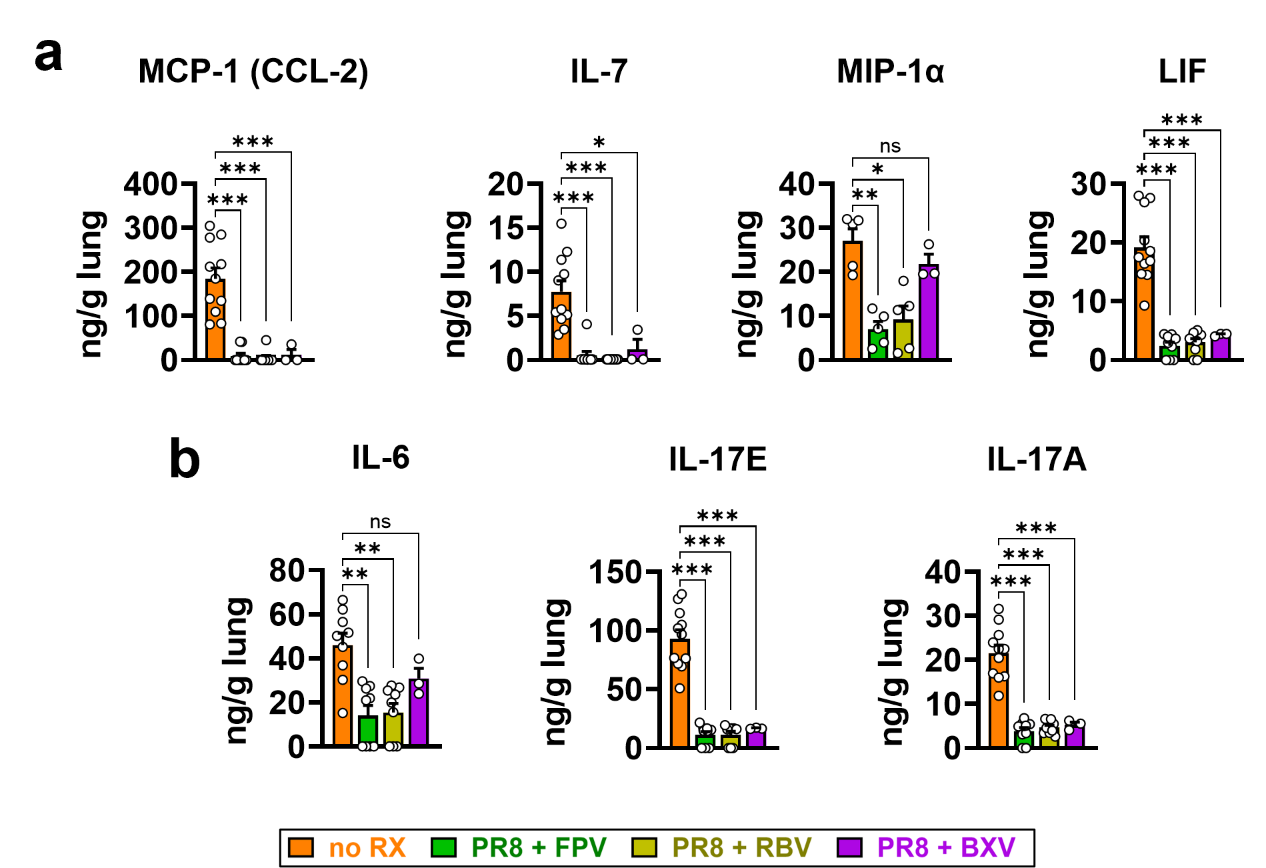

**Fig. S8 | Expression of selected cytokines and chemokines in influenza-infected lungs following antiviral therapy.** **a,** Neutrophil-mediated inflammatory mediators MCP-1, IL-7, MIP-1α, and LIF; and **b**, systemic inflammatory mediators IL-6, IL-17E, and Il-17A expression in lung tissues of sham-treated animals (no Rx) during peak disease at 6 days post-infection (dpi) or at 9 dpi in mice given antiviral treatment with either favipiravir (FPV), ribavirin (RBV), or baloxavir marboxil (BXV). Data are mean ± s.d. Statistical analysis was performed using Welch’s ANOVA with Dunnett’s correction; * *p* < 0.05; ** *p* < 0.005; *** *p* < 0.001; *ns*, not significant.

**Figure S9**

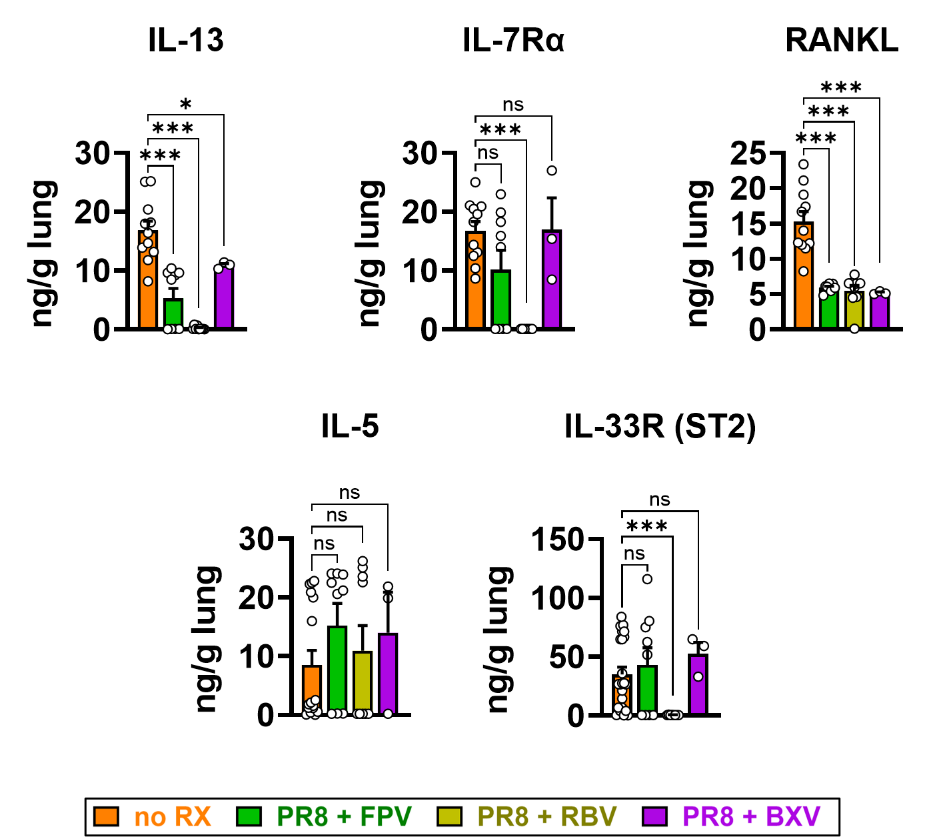

**Fig. S9 | Expression of selected cytokines and chemokines that are distinctly expressed in the lungs of mice treated with ribavirin (RBV).** Expression levels of IL-13, IL-7Rα, RANKL, IL-5, and IL-3R (ST2) in lung tissues of sham-treated animals (no Rx) during peak disease at 6 days post-infection (dpi) or at 9 dpi in mice given antiviral treatment with either favipiravir (FPV), ribavirin (RBV), or baloxavir marboxil (BXV). Data are mean ± s.d. Statistical analysis was performed using Welch’s ANOVA; * *p* < 0.05; ** *p* < 0.005; *** *p* < 0.001; *ns*, not significant.

**Figure S10**

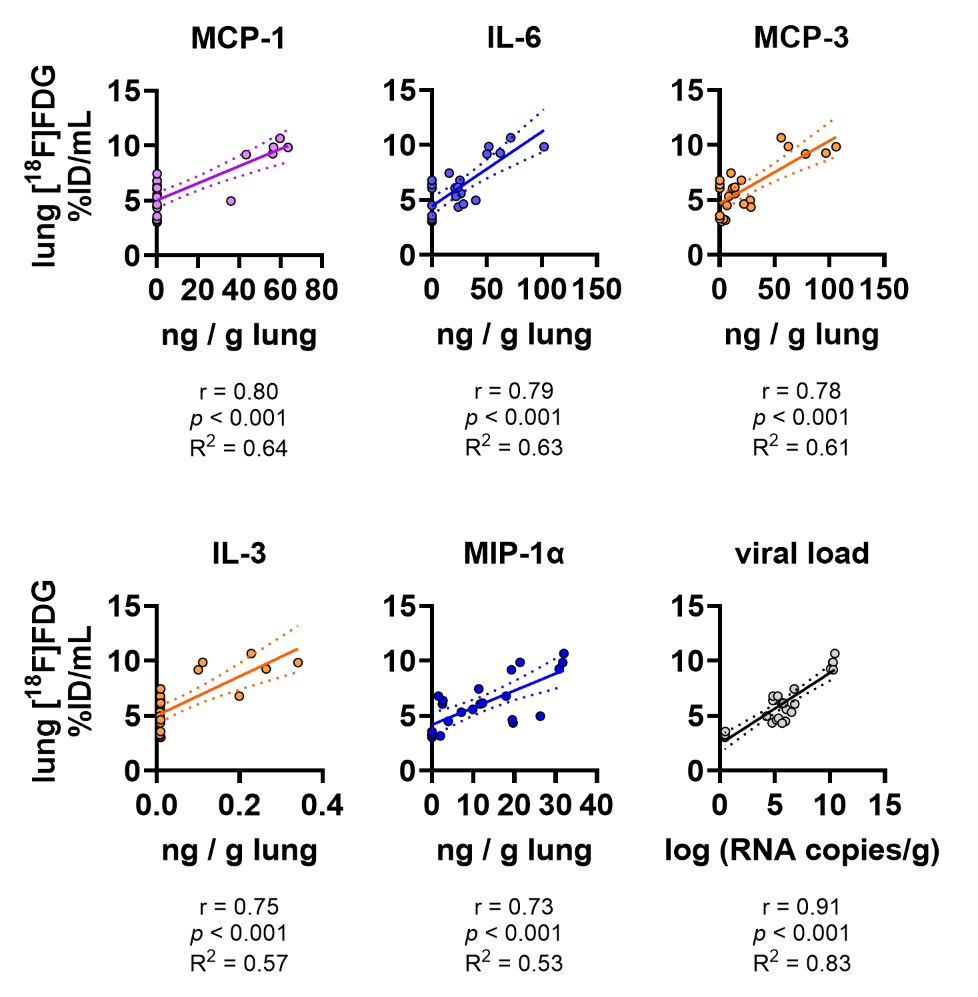

**Fig. S10 | Linear correlation between lung [^18^F]FDG uptake and measures of infection and inflammation in a mouse model of influenza A/PR8 infection.** Scatter plots depicting linear relationships between lung [¹⁸F]FDG uptake and parameters with the strongest correlations: MCP-1, LIF, MIP-1α, IFN-α, IL-6 and viral load in the lungs. *r*, Pearson coefficient. *R^2^*, linearity coefficient.

**Figure S11**

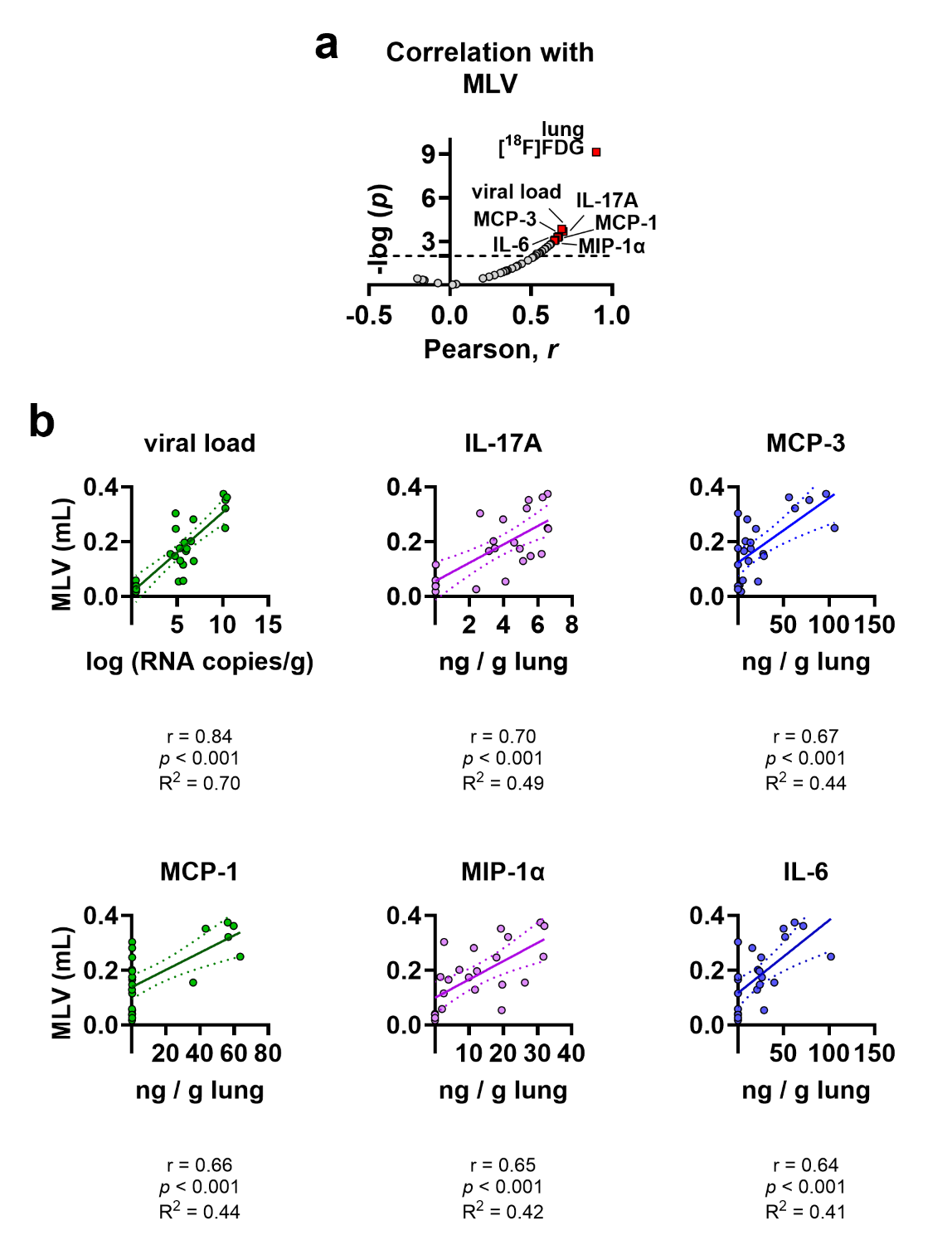

**Fig. S11 | Linear association between metabolic lung volume (MLV) and measures of infection and inflammation in a mouse model of influenza A/PR8 infection. a,** Volcano plot depicting correlations between MLV and infection and inflammation parameters. **b**, Scatter plots depicting linear relationships between lung MLV and parameters with the strongest correlations: lung viral load, and expression of cytokines: IL17-A, MCP-3, MCP-1, MIP-1α, and IL-6. *r*, Pearson coefficient. *R^2^*, linearity coefficient.

**Figure S12**

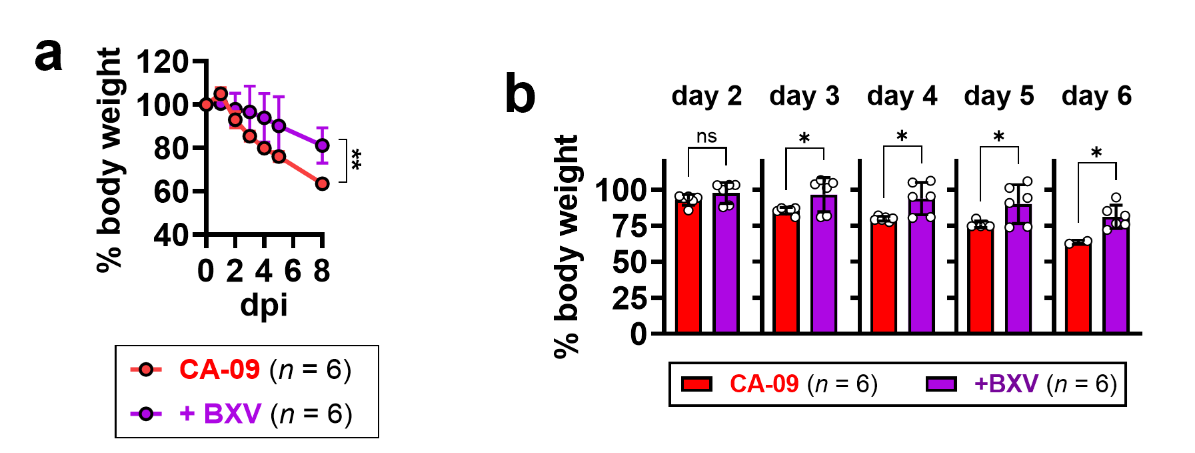

**Fig. S12 | Changes in body weight over time in mice infected with H1N1/CA-09. a,** Body weights, expressed as % relative to pre-infection, following infection and treatment with the antiviral baloxavir marboxil (BXV). **b**, Comparison of body weights on days 2-6 post-infection (dpi). Data are shown as mean ± sd of 6 mice per group representative of 3 independent experiments. Means of AUCs of body weight trend in ***a*** and daily body weights in ***b*** were compared by Welch’s *t*-test. * *p* < 0.05; ** *p* < 0.005; *** *p* < 0.001; *ns*, not significant.

**Figure S13**

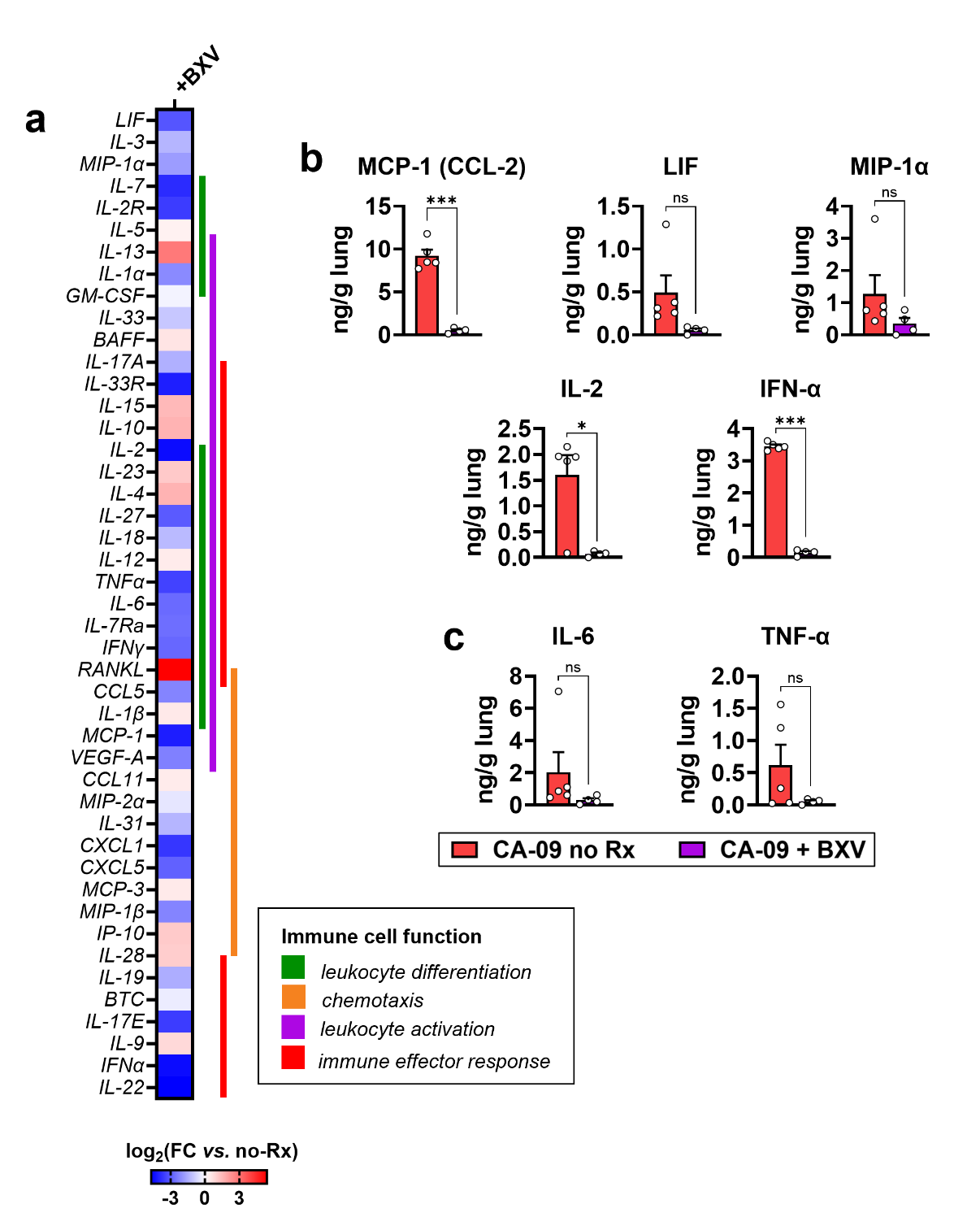

**Fig. S13 | Lung inflammation in mice infected with H1N1/CA-09. a,** Heatmap of selected cytokines and chemokines involved in immune cells differentiation, activation, chemotaxis, and effector response in the lungs of infected mice treated with baloxavir marboxil (BXV). Data are shown as log_2_ fold-change expression relative to vehicle-treated mouse lungs (log₂FC *vs.* no-Rx). Proteins associated with leukocyte activation (*purple*), differentiation (*green*), or both (*blue*) are indicated. Proteins involved in immune effector response are shown in *red*. **b-c,** Expression of selected cytokines and chemokines in influenza-infected lungs following BXV therapy: **b,** Neutrophil-mediated inflammatory mediators MCP-1, LIF, MIP-1α, IL-2, and IFN-α; and **c**, systemic inflammatory mediators IL-6 and TNF-α expression in lung tissues of CA-09 infected, sham-treated animals (no Rx) during peak disease at 6 days post-infection (dpi) or at 9 dpi in mice given BXV treatment. Data are presented as mean ± sd of 6 mice per group, representative of 3 independent experiments. Statistical analysis was performed using Welch’s *t*-test; * *p* < 0.05; ** *p* < 0.005; *** *p* < 0.001; *ns*, not significant.

**Figure S14**

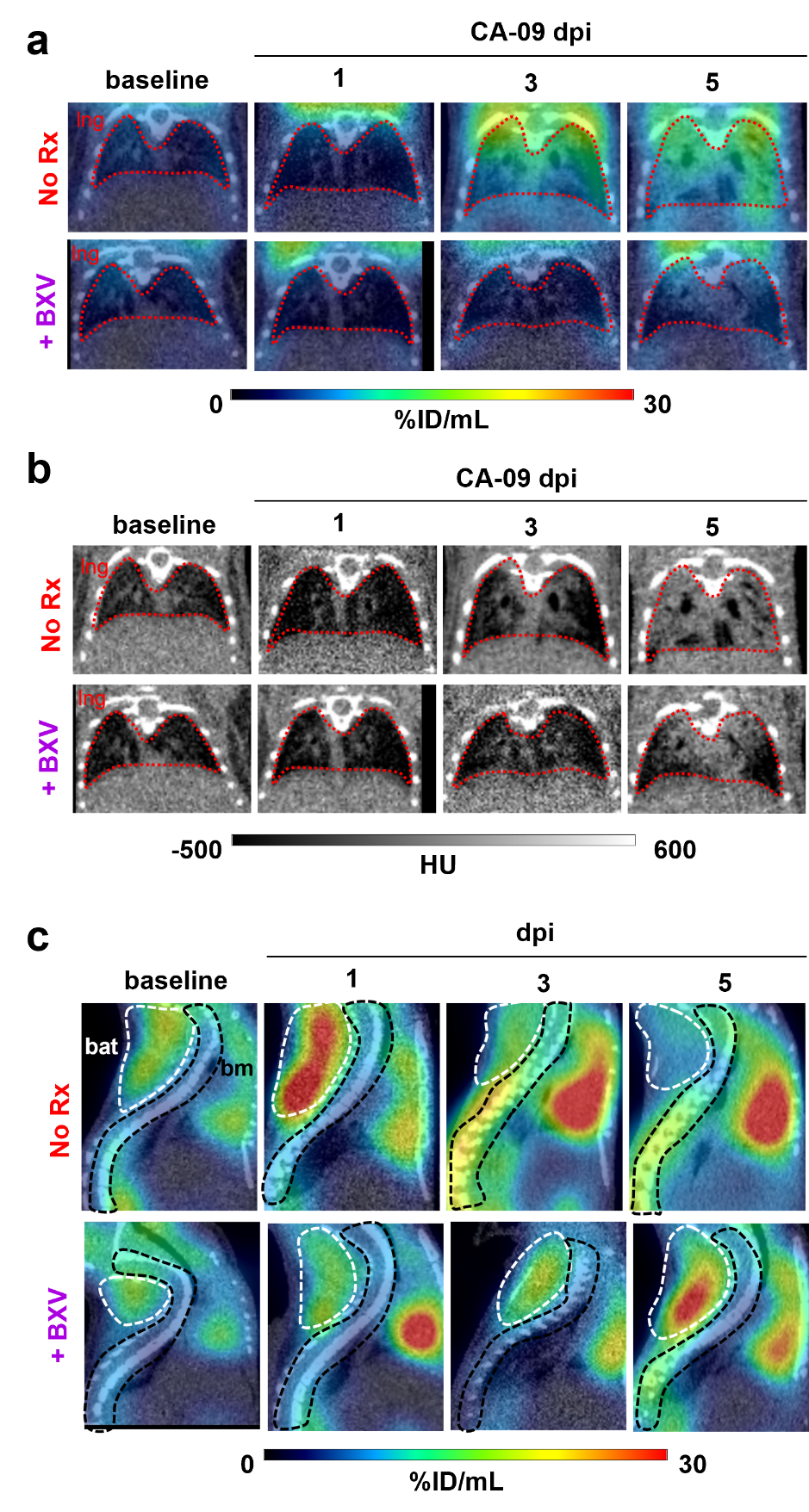

**Fig. S14 |Dynamics of [^18^F]FDG uptake in CA-09-infected lungs revealed in PET scans. a,b,** Representative transverse slices of **a,** PET/CT and **b,** CT images acquired at indicated days post-infection (dpi) and focused on the lung region. **c,** Representative sagittal views of the thoracic cavity showing [¹⁸F]FDG uptake in lung, brown adipose tissue, and bone marrow. Volumes of interest (VOIs) representing lungs (*lng*) are indicated by red dashed lines; brown adipose tissue (*bat*) by white dashed lines; and bone marrow (*bm*) by black dashed lines. PET images are displayed at 0–30% ID/mL, and CT images at −500 to 600 Hounsfield units (HU), as indicated by the colour scale bars.

**Figure S15**

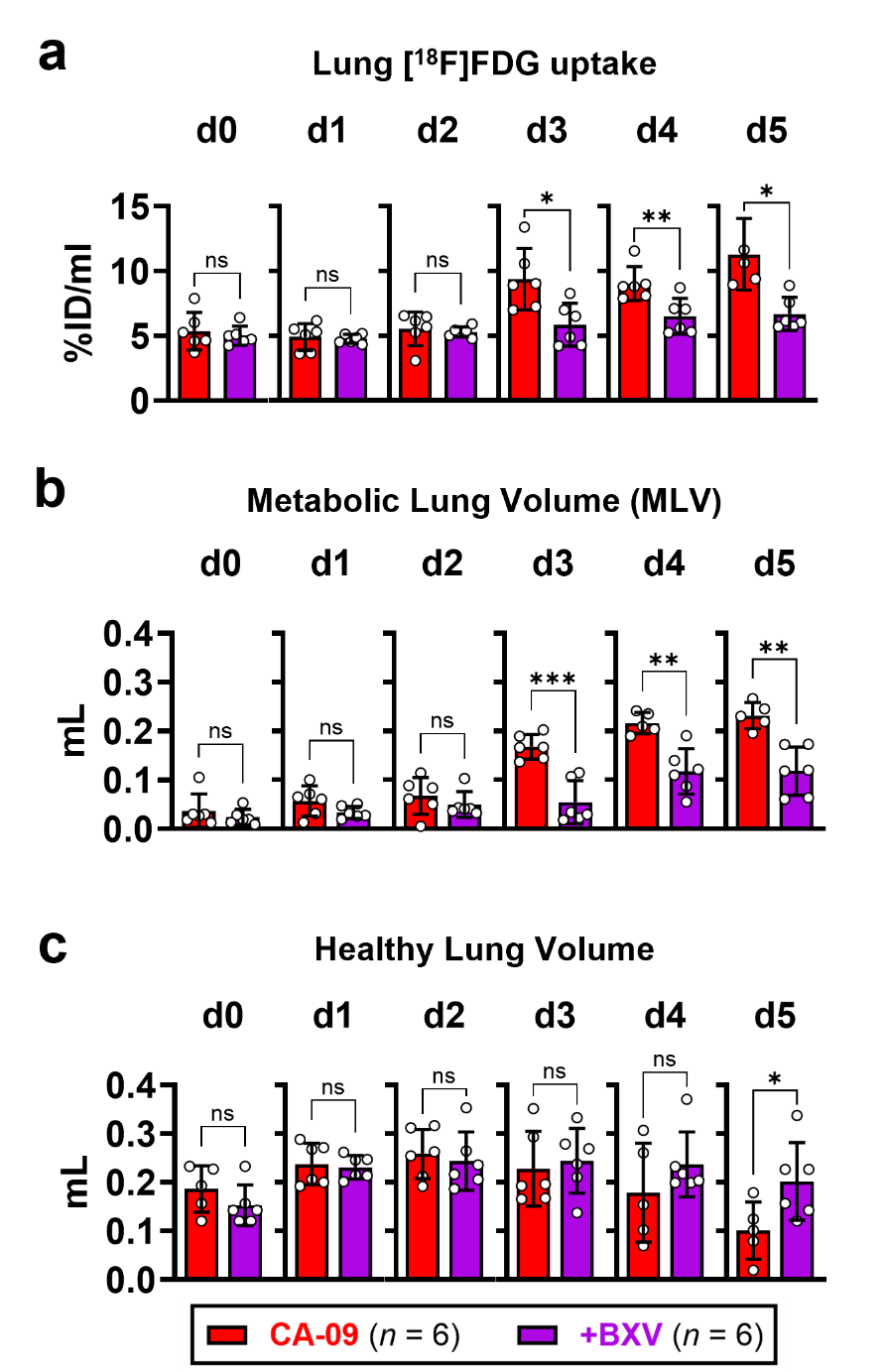

**Fig. S15 | Differentiation between antiviral treatment and control treatment in lungs from mice infected with CA-09-infected. a,** Lung [^18^F]FDG uptake values; **b**, metabolic lung volumes (MLV); and **c**, healthy lung volumes between mice treated with baloxavir marboxil (BXV) and vehicle on various days post-infection. Data are mean ± sd from 6 mice per group, representative of 3 independent experiments. Statistical analysis was performed using Welch’s *t*-test; * *p* < 0.05; ** *p* < 0.005; *ns*, not significant.

**Figure S16**

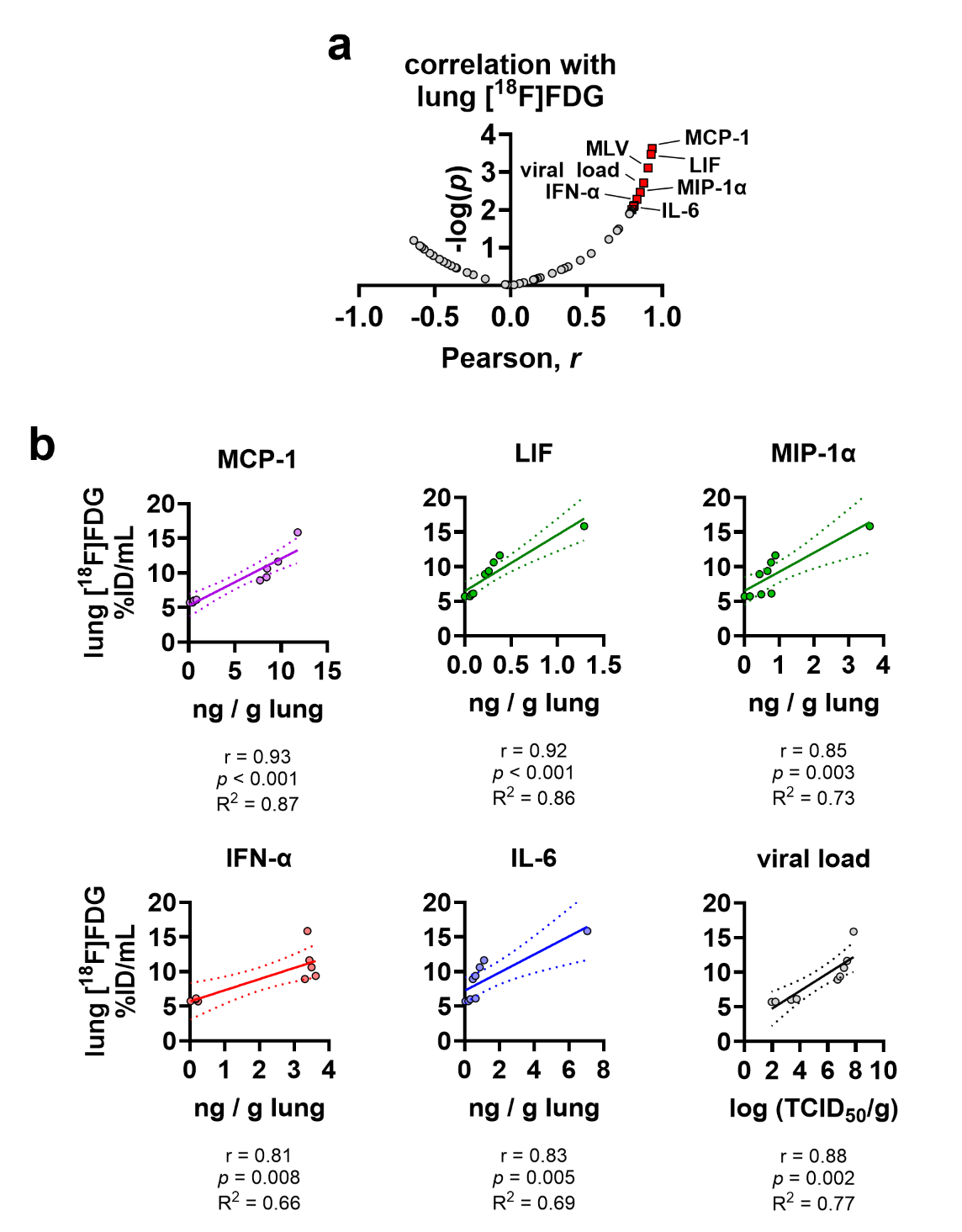

**Fig. S16 | Linear association between lung [^18^F]FDG uptake and measures of infection and inflammation in a mouse model of influenza A/CA-09 infection. a,** Volcano plot depicting correlations between lung [^18^F]FDG uptake and infection and inflammation parameters. **b**, Scatter plots depicting linear relationships between lung [^18^F]FDG uptake and parameters with the strongest correlations: MCP-1, LIF, MIP-1α, IFN- α, IL-6, and lung viral load. *r*, Pearson coefficient. *R^2^*, linearity coefficient.

**Figure S17**

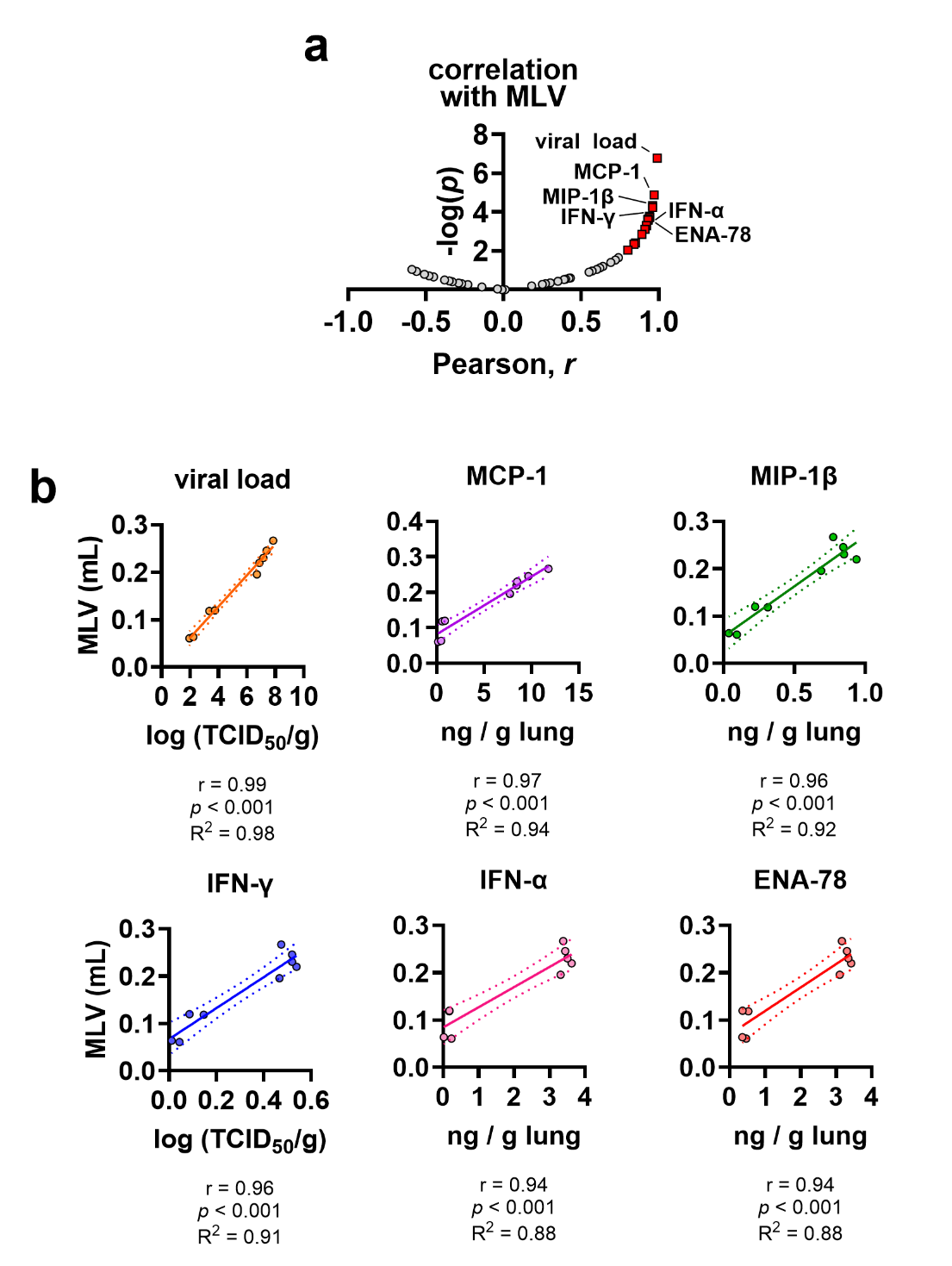

**Fig. S17 | Linear association between MLV and measures of infection and inflammation in a mouse model of influenza A/CA-09 infection. a,** Volcano plot depicting correlations between MLV and infection and inflammation parameters. **b**, Scatter plots depicting linear relationships between MLV and parameters with the strongest correlations: MCP-1, LIF, MIP-1α, IFN- α, IL-6, and lung viral load. *r*, Pearson coefficient. *R^2^*, linearity coefficient.

**Figure S18**

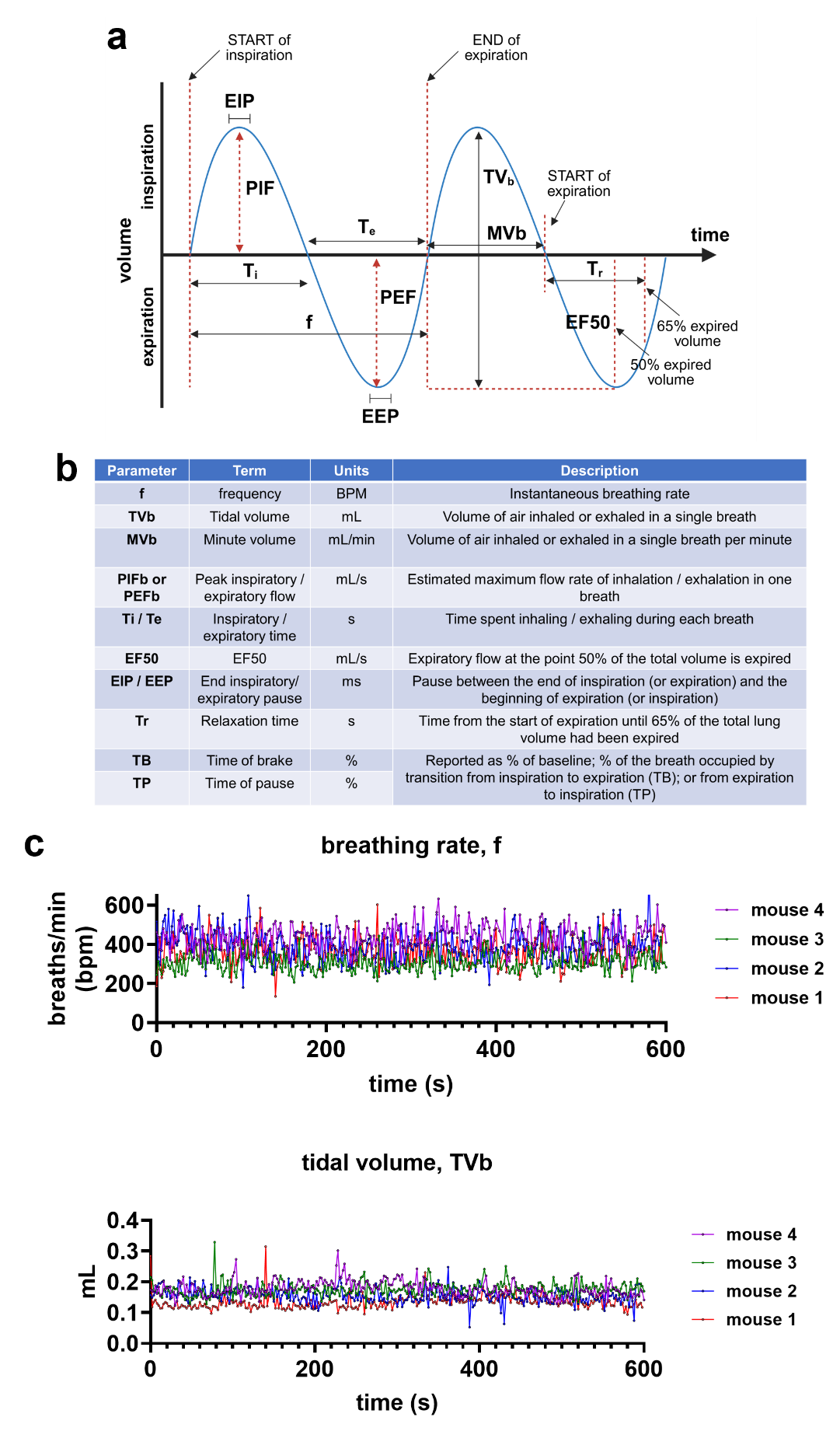

**Fig. S18 | Respiratory parameters measured by whole-body plethysmography.**
**a,** Schematic representation of the respiratory cycle, showing changes in chamber volume over time. **b,** Table defining the respiratory parameters quantified by whole-body plethysmography (WBP). **c,** Sample plot depicting breathing rate and tidal volume recorded over 10 minutes.

**Figure S19**

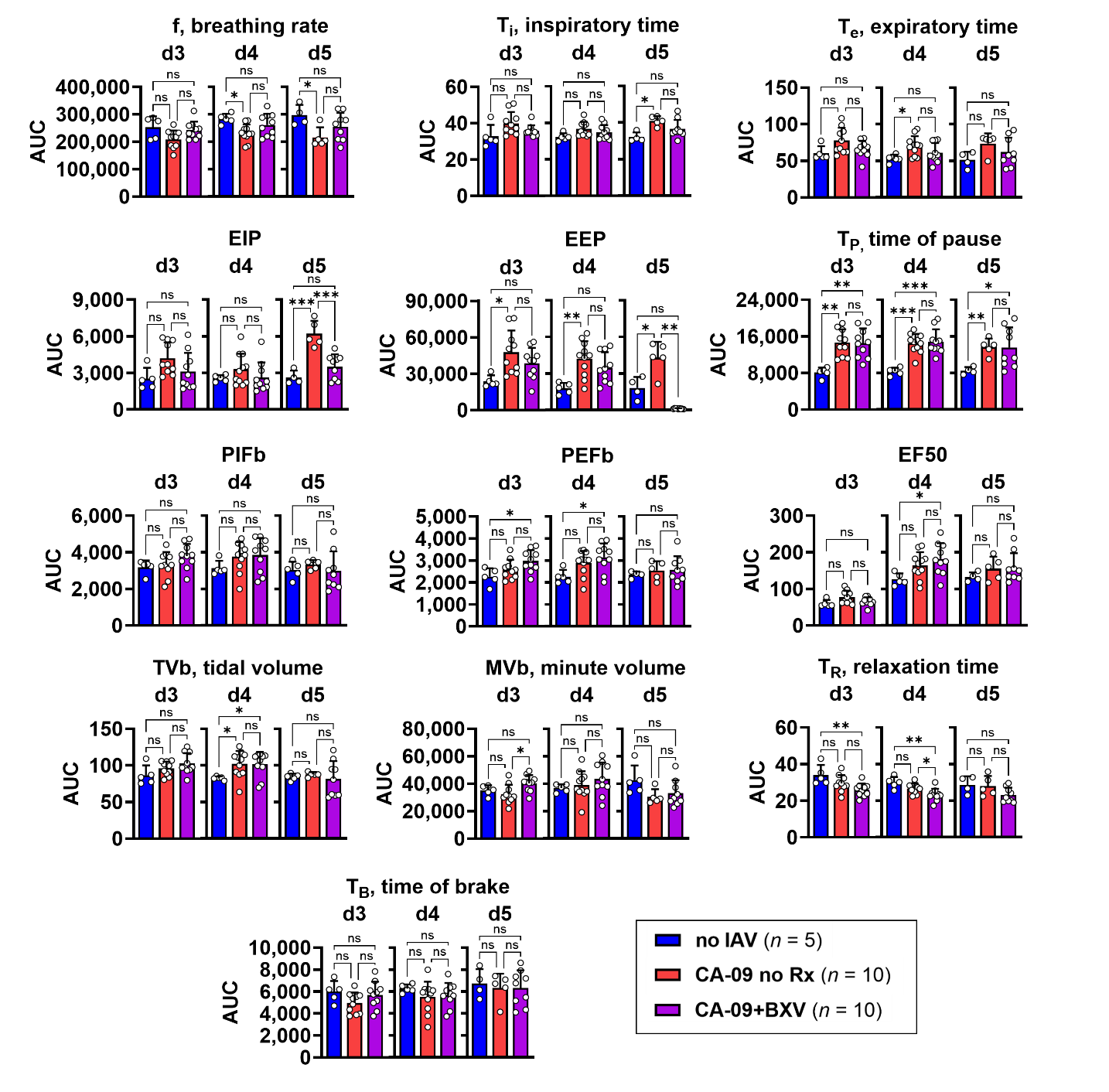

**Fig. S19 | Area under the curve analysis of respiratory parameters measured by whole-body plethysmography.** Mice were infected with influenza A/California/04/09 (CA-09) and treated with baloxavir marboxil (BXV) or vehicle control (no Rx). Sham-infected mice treated with saline (no IAV) served as controls. Respiratory parameters were recorded continuously by whole-body plethysmography (WBP) every 2 s over a 10-min period at 3, 4, and 5 days post-infection (dpi). Data are shown as area under the curve (AUC) values for each recording period and presented as mean ± s.d. Statistical significance was assessed using Welch’s ANOVA; * *p* < 0.05; ** *p* < 0.005; *** *p* < 0.001; ns, not significant.

**Supplementary Tables**

**Table S1 | Performance of lung [^18^F]FDG uptake reported as %ID/mL at 3 days post-infection (dpi) with PR8 strain in detecting influenza-associated lung inflammation and predicting lethality by 6 dpi**. Excerpt of model specificity and sensitivity data obtained from receiving operator characteristic (ROC) analysis in GraphPad Prism v11.0.2. Negative predictive values (NPV) and positive predictive values (PPV) were calculated based on an assumption of high lethality rate (95%) of influenza infection.

| Cut-off | Sensitivity% | 95% CI | Specificity% | 95% CI | Likelihood ratio | NPV_high_ | PPV_high_ |
| --- | --- | --- | --- | --- | --- | --- | --- |
| > 3.050 | 100 | 83.18% to 100.0% | 4 | 1.566% to 9.837% | 1.042 | 1.00 | 0.95 |
| > 3.345 | 100 | 83.18% to 100.0% | 9 | 4.807% to 16.23% | 1.099 | 1.00 | 0.95 |
| > 3.605 | 100 | 83.18% to 100.0% | 16 | 10.10% to 24.42% | 1.19 | 1.00 | 0.96 |
| > 3.710 | 100 | 83.18% to 100.0% | 23 | 15.84% to 32.15% | 1.299 | 1.00 | 0.96 |
| > 3.855 | 100 | 83.18% to 100.0% | 30 | 21.89% to 39.58% | 1.429 | 1.00 | 0.96 |
| > 3.940 | 100 | 83.18% to 100.0% | 35 | 26.36% to 44.75% | 1.538 | 1.00 | 0.97 |
| > 4.045 | 100 | 83.18% to 100.0% | 40 | 30.94% to 49.80% | 1.667 | 1.00 | 0.97 |
| > 4.190 | 100 | 83.18% to 100.0% | 45 | 35.61% to 54.76% | 1.818 | 1.00 | 0.97 |
| > 4.365 | 100 | 83.18% to 100.0% | 56 | 46.23% to 65.33% | 2.273 | 1.00 | 0.98 |
| > 4.560 | 100 | 83.18% to 100.0% | 63 | 53.22% to 71.82% | 2.703 | 1.00 | 0.98 |
| > 4.685 | 100 | 83.18% to 100.0% | 71 | 61.46% to 78.99% | 3.448 | 1.00 | 0.98 |
| > 4.865 | 94.74 | 75.36% to 99.73% | 77 | 67.85% to 84.16% | 4.119 | 0.44 | 0.99 |
| > 5.080 | 94.74 | 75.36% to 99.73% | 84 | 75.58% to 89.90% | 5.921 | 0.46 | 0.99 |
| > 5.450 | 89.47 | 68.61% to 98.13% | 89 | 81.37% to 93.75% | 8.134 | 0.31 | 0.99 |
| > 5.870 | 84.21 | 62.43% to 94.48% | 93 | 86.25% to 96.57% | 12.03 | 0.24 | 1.00 |
| > 6.130 | 68.42 | 46.01% to 84.64% | 97 | 91.55% to 99.18% | 22.81 | 0.14 | 1.00 |
| > 6.755 | 42.11 | 23.14% to 63.72% | 98 | 93.00% to 99.64% | 21.05 | 0.08 | 1.00 |

**Table S2 | Performance of lung [^18^F]FDG uptake reported as %ID/mL at 6 days post-infection (dpi) with PR8 strain in detecting influenza-associated lung inflammation and predicting lethality by 6 dpi** . Excerpt of model specificity and sensitivity data obtained from receiving operator characteristic (ROC) analysis in GraphPad Prism v11.0.2. Negative predictive values (NPV) and positive predictive values (PPV) were calculated based on an assumption of high lethality rate (95%) of influenza infection.

| Cut-off | Sensitivity% | 95% CI | Specificity% | 95% CI | Likelihood ratio | NPV_high_ | PPV_high_ |
| --- | --- | --- | --- | --- | --- | --- | --- |
| > 3.050 | 100 | 87.54% to 100.0% | 4 | 1.566% to 9.837% | 1.042 | 1.00 | 0.95 |
| > 3.345 | 100 | 87.54% to 100.0% | 9 | 4.807% to 16.23% | 1.099 | 1.00 | 0.95 |
| > 3.605 | 100 | 87.54% to 100.0% | 16 | 10.10% to 24.42% | 1.19 | 1.00 | 0.96 |
| > 3.710 | 100 | 87.54% to 100.0% | 23 | 15.84% to 32.15% | 1.299 | 1.00 | 0.96 |
| > 3.855 | 100 | 87.54% to 100.0% | 30 | 21.89% to 39.58% | 1.429 | 1.00 | 0.96 |
| > 3.940 | 100 | 87.54% to 100.0% | 35 | 26.36% to 44.75% | 1.538 | 1.00 | 0.97 |
| > 4.045 | 100 | 87.54% to 100.0% | 40 | 30.94% to 49.80% | 1.667 | 1.00 | 0.97 |
| > 4.190 | 100 | 87.54% to 100.0% | 45 | 35.61% to 54.76% | 1.818 | 1.00 | 0.97 |
| > 4.365 | 100 | 87.54% to 100.0% | 56 | 46.23% to 65.33% | 2.273 | 1.00 | 0.98 |
| > 4.560 | 100 | 87.54% to 100.0% | 63 | 53.22% to 71.82% | 2.703 | 1.00 | 0.98 |
| > 4.685 | 100 | 87.54% to 100.0% | 71 | 61.46% to 78.99% | 3.448 | 1.00 | 0.98 |
| > 4.935 | 100 | 87.54% to 100.0% | 78 | 68.93% to 85.00% | 4.545 | 1.00 | 0.99 |
| > 5.180 | 100 | 87.54% to 100.0% | 85 | 76.72% to 90.69% | 6.667 | 1.00 | 0.99 |
| > 5.470 | 96.3 | 81.72% to 99.81% | 89 | 81.37% to 93.75% | 8.754 | 0.56 | 0.99 |
| > 5.995 | 92.59 | 76.63% to 98.68% | 95 | 88.82% to 97.85% | 18.52 | 0.40 | 1.00 |
| > 7.290 | 85.19 | 67.52% to 94.08% | 99 | 94.55% to 99.95% | 85.19 | 0.26 | 1.00 |

**Table S3 | Performance of lung [^18^F]FDG uptake reported as %ID/mL at 3 days post-infection (dpi) with PR8 strain in detecting dampened influenza-associated lung inflammation and predicting survival beyond 6 dpi**. Excerpt of model specificity and sensitivity data obtained from receiving operator characteristic (ROC) analysis in GraphPad Prism v11.0.2. Two sets of negative predictive values (NPV) and positive predictive values (PPV) were calculated based on assumptions of low survival rate (50%) or high survival rate (95%) following therapeutic intervention.

| Cut-off | Sensitivity% | 95% CI | Specificity% | 95% CI | Likelihood ratio | NPV_low_*^a^* | PPV_low_*^b^* | NPV_high_*^c^* | PPV_high_*^d^* |
| --- | --- | --- | --- | --- | --- | --- | --- | --- | --- |
| < 3.510 | 15.38 | 2.734% to 42.23% | 100 | 83.18% to 100.0% |  | 0.54 | 1.00 | 0.23 | 1.00 |
| < 4.050 | 38.46 | 17.71% to 64.48% | 100 | 83.18% to 100.0% |  | 0.62 | 1.00 | 0.29 | 1.00 |
| < 4.935 | 53.85 | 29.14% to 76.79% | 94.74 | 75.36% to 99.73% | 10.23 | 0.67 | 0.91 | 0.34 | 0.98 |
| < 5.160 | 76.92 | 49.74% to 91.82% | 94.74 | 75.36% to 99.73% | 14.62 | 0.80 | 0.94 | 0.51 | 0.98 |
| < 5.535 | 92.31 | 66.69% to 99.61% | 89.47 | 68.61% to 98.13% | 8.769 | 0.92 | 0.90 | 0.74 | 0.97 |
| < 6.070 | 92.31 | 66.69% to 99.61% | 73.68 | 51.21% to 88.19% | 3.508 | 0.91 | 0.78 | 0.71 | 0.93 |
| < 6.330 | 92.31 | 66.69% to 99.61% | 57.89 | 36.28% to 76.86% | 2.192 | 0.88 | 0.69 | 0.65 | 0.90 |
| < 6.755 | 100 | 77.19% to 100.0% | 42.11 | 23.14% to 63.72% | 1.727 | 1.00 | 0.63 | 1.00 | 0.87 |
| < 7.870 | 100 | 77.19% to 100.0% | 26.32 | 11.81% to 48.79% | 1.357 | 1.00 | 0.58 | 1.00 | 0.84 |
| < 8.955 | 100 | 77.19% to 100.0% | 10.53 | 1.870% to 31.39% | 1.118 | 1.00 | 0.53 | 1.00 | 0.82 |

***^a^*** NPV when treatment-induced survival rate is low (50%)

***^b^*** PPV when treatment-induced survival rate is low (50%)

***^c^*** NPV when treatment-induced survival rate is high (95%)

***^d^*** PPV when treatment-induced survival rate is high (95%)

**Table S4 | Performance of lung [^18^F]FDG uptake reported as %ID/mL at 6 days post-infection (dpi) with PR8 strain in detecting dampened influenza-associated lung inflammation and predicting survival beyond 6 dpi**. Excerpt of model specificity and sensitivity data obtained from receiving operator characteristic (ROC) analysis in GraphPad Prism v11.0.2. Two sets of negative predictive values (NPV) and positive predictive values (PPV) were calculated based on assumptions of low survival rate (50%) or high survival rate (95%) following therapeutic intervention.

| Cut-off | Sensitivity% | 95% CI | Specificity% | 95% CI | Likelihood ratio | NPV_low_*^a^* | PPV_low_*^b^* | NPV_high_*^c^* | PPV_high_*^d^* |
| --- | --- | --- | --- | --- | --- | --- | --- | --- | --- |
| < 4.415 | 8.333 | 1.481% to 25.85% | 100 | 87.54% to 100.0% |  | 0.52 | 1.00 | 0.21 | 1.00 |
| < 4.780 | 20.83 | 9.245% to 40.47% | 100 | 87.54% to 100.0% |  | 0.56 | 1.00 | 0.24 | 1.00 |
| < 5.130 | 33.33 | 17.97% to 53.29% | 100 | 87.54% to 100.0% |  | 0.60 | 1.00 | 0.27 | 1.00 |
| < 5.635 | 41.67 | 24.47% to 61.17% | 96.3 | 81.72% to 99.81% | 11.25 | 0.62 | 0.92 | 0.29 | 0.98 |
| < 5.910 | 54.17 | 35.07% to 72.11% | 96.3 | 81.72% to 99.81% | 14.63 | 0.68 | 0.94 | 0.34 | 0.98 |
| < 6.125 | 66.67 | 46.71% to 82.03% | 92.59 | 76.63% to 98.68% | 9 | 0.74 | 0.90 | 0.41 | 0.97 |
| < 6.350 | 79.17 | 59.53% to 90.76% | 85.19 | 67.52% to 94.08% | 5.344 | 0.80 | 0.84 | 0.51 | 0.96 |
| < 6.980 | 91.67 | 74.15% to 98.52% | 85.19 | 67.52% to 94.08% | 6.188 | 0.91 | 0.86 | 0.72 | 0.96 |
| < 7.535 | 100 | 86.20% to 100.0% | 81.48 | 63.30% to 91.82% | 5.4 | 1.00 | 0.84 | 1.00 | 0.96 |
| < 7.965 | 100 | 86.20% to 100.0% | 70.37 | 51.52% to 84.15% | 3.375 | 1.00 | 0.77 | 1.00 | 0.93 |
| < 8.740 | 100 | 86.20% to 100.0% | 51.85 | 33.99% to 69.26% | 2.077 | 1.00 | 0.67 | 1.00 | 0.89 |
| < 9.705 | 100 | 86.20% to 100.0% | 33.33 | 18.64% to 52.18% | 1.5 | 1.00 | 0.60 | 1.00 | 0.86 |
| < 10.61 | 100 | 86.20% to 100.0% | 14.81 | 5.916% to 32.48% | 1.174 | 1.00 | 0.54 | 1.00 | 0.82 |

***^a^*** NPV when treatment-induced survival rate is low (50%)

***^b^*** PPV when treatment-induced survival rate is low (50%)

***^c^*** NPV when treatment-induced survival rate is high (95%)

***^d^*** PPV when treatment-induced survival rate is high (95%)

**Table S5 | Performance of metabolic lung volumes (MLV) at 3 days post-infection (dpi) with PR8 strain in detecting influenza-associated lung inflammation and predicting lethality by 6 dpi**. Excerpt of model specificity and sensitivity data obtained from receiving operator characteristic (ROC) analysis in GraphPad Prism v11.0.2. Negative predictive values (NPV) and positive predictive values (PPV) were calculated based on an assumption of high lethality rate (95%) of influenza infection.

| Cut-off | Sensitivity% | 95% CI | Specificity% | 95% CI | Likelihood ratio | NPV_high_ | PPV_high_ |
| --- | --- | --- | --- | --- | --- | --- | --- |
| > 0.02500 | 100 | 85.13% to 100.0% | 28.13 | 18.59% to 40.13% | 1.391 | 1.00 | 0.96 |
| > 0.06500 | 100 | 85.13% to 100.0% | 73.44 | 61.52% to 82.70% | 3.765 | 1.00 | 0.99 |
| > 0.07500 | 100 | 85.13% to 100.0% | 79.69 | 68.29% to 87.73% | 4.923 | 1.00 | 0.99 |
| > 0.08500 | 95.45 | 78.20% to 99.77% | 84.38 | 73.57% to 91.29% | 6.109 | 0.49 | 0.99 |
| > 0.1050 | 95.45 | 78.20% to 99.77% | 90.63 | 81.02% to 95.63% | 10.18 | 0.51 | 0.99 |
| > 0.1350 | 95.45 | 78.20% to 99.77% | 96.88 | 89.30% to 99.44% | 30.55 | 0.53 | 1.00 |
| > 0.1550 | 86.36 | 66.67% to 95.25% | 98.44 | 91.67% to 99.92% | 55.27 | 0.28 | 1.00 |
| > 0.2050 | 54.55 | 34.66% to 73.08% | 100 | 94.34% to 100.0% |  | 0.10 | 1.00 |
| > 0.2650 | 31.82 | 16.36% to 52.68% | 100 | 94.34% to 100.0% |  | 0.07 | 1.00 |
| > 0.3450 | 9.091 | 1.615% to 27.81% | 100 | 94.34% to 100.0% |  | 0.05 | 1.00 |

**Table S6 | Performance of metabolic lung volumes (MLV) at 6 days post-infection (dpi) with PR8 strain in detecting influenza-associated lung inflammation and predicting lethality by 6 dpi**. Excerpt of model specificity and sensitivity data obtained from receiving operator characteristic (ROC) analysis in GraphPad Prism v11.0.2. Negative predictive values (NPV) and positive predictive values (PPV) were calculated based on an assumption of high lethality rate (95%) of influenza infection.

| Cut-off | Sensitivity% | 95% CI | Specificity% | 95% CI | Likelihood ratio | NPV_high_ | PPV_high_ |
| --- | --- | --- | --- | --- | --- | --- | --- |
| > 0.01500 | 100 | 90.11% to 100.0% | 14.06 | 7.579% to 24.62% | 1.164 | 1.00 | 0.96 |
| > 0.02500 | 100 | 90.11% to 100.0% | 28.13 | 18.59% to 40.13% | 1.391 | 1.00 | 0.96 |
| > 0.03500 | 100 | 90.11% to 100.0% | 48.44 | 36.63% to 60.42% | 1.939 | 1.00 | 0.97 |
| > 0.04500 | 100 | 90.11% to 100.0% | 62.5 | 50.25% to 73.33% | 2.667 | 1.00 | 0.98 |
| > 0.05500 | 97.14 | 85.47% to 99.85% | 70.31 | 58.23% to 80.09% | 3.272 | 0.56 | 0.98 |
| > 0.07500 | 97.14 | 85.47% to 99.85% | 79.69 | 68.29% to 87.73% | 4.782 | 0.59 | 0.99 |
| > 0.1050 | 91.43 | 77.62% to 97.04% | 90.63 | 81.02% to 95.63% | 9.752 | 0.36 | 0.99 |
| > 0.1450 | 82.86 | 67.32% to 91.90% | 96.88 | 89.30% to 99.44% | 26.51 | 0.23 | 1.00 |
| > 0.1750 | 71.43 | 54.95% to 83.67% | 100 | 94.34% to 100.0% |  | 0.16 | 1.00 |

**Table S7 | Performance of metabolic lung volumes (MLV) at 3 days post-infection (dpi) with PR8 strain in detecting dampened influenza-associated lung inflammation and predicting survival beyond 6 dpi**. Excerpt of model specificity and sensitivity data obtained from receiving operator characteristic (ROC) analysis in GraphPad Prism v11.0.2. Two sets of negative predictive values (NPV) and positive predictive values (PPV) were calculated based on assumptions of low survival rate (50%) or high survival rate (95%) following therapeutic intervention.

| Cut-off | Sensitivity% | 95% CI | Specificity% | 95% CI | Likelihood ratio | NPV_low_*^a^* | PPV_low_*^b^* | NPV_high_*^c^* | PPV_high_*^d^* |
| --- | --- | --- | --- | --- | --- | --- | --- | --- | --- |
| < 0.06000 | 20.83 | 9.245% to 40.47% | 100 | 85.13% to 100.0% |  | 0.56 | 1.00 | 0.24 | 1.00 |
| < 0.1000 | 50 | 31.43% to 68.57% | 95.45 | 78.20% to 99.77% | 11 | 0.66 | 0.92 | 0.32 | 0.98 |
| < 0.1350 | 62.5 | 42.71% to 78.84% | 95.45 | 78.20% to 99.77% | 13.75 | 0.72 | 0.93 | 0.39 | 0.98 |
| < 0.1650 | 87.5 | 69.00% to 95.66% | 72.73 | 51.85% to 86.85% | 3.208 | 0.85 | 0.76 | 0.59 | 0.93 |
| < 0.1750 | 87.5 | 69.00% to 95.66% | 63.64 | 42.95% to 80.27% | 2.406 | 0.84 | 0.71 | 0.56 | 0.91 |
| < 0.1850 | 95.83 | 79.76% to 99.79% | 63.64 | 42.95% to 80.27% | 2.635 | 0.94 | 0.72 | 0.79 | 0.91 |
| < 0.1950 | 95.83 | 79.76% to 99.79% | 59.09 | 38.73% to 76.74% | 2.343 | 0.93 | 0.70 | 0.78 | 0.90 |
| < 0.2050 | 95.83 | 79.76% to 99.79% | 54.55 | 34.66% to 73.08% | 2.108 | 0.93 | 0.68 | 0.77 | 0.89 |
| < 0.2250 | 95.83 | 79.76% to 99.79% | 40.91 | 23.26% to 61.27% | 1.622 | 0.91 | 0.62 | 0.71 | 0.87 |
| < 0.2650 | 100 | 86.20% to 100.0% | 31.82 | 16.36% to 52.68% | 1.467 | 1.00 | 0.59 | 1.00 | 0.85 |

***^a^*** NPV when treatment-induced survival rate is low (50%)

***^b^*** PPV when treatment-induced survival rate is low (50%)

***^c^*** NPV when treatment-induced survival rate is high (95%)

***^d^*** PPV when treatment-induced survival rate is high (95%)

**Table S8 | Performance of metabolic lung volumes (MLV) at 6 days post-infection (dpi) with PR8 strain in detecting dampened influenza-associated lung inflammation and predicting survival beyond 6 dpi**. Excerpt of model specificity and sensitivity data obtained from receiving operator characteristic (ROC) analysis in GraphPad Prism v11.0.2. Two sets of negative predictive values (NPV) and positive predictive values (PPV) were calculated based on assumptions of low survival rate (50%) or high survival rate (95%) following therapeutic intervention.

| Cut-off | Sensitivity% | 95% CI | Specificity% | 95% CI | Likelihood ratio | NPV_low_*^a^* | PPV_low_*^b^* | NPV_high_*^c^* | PPV_high_*^d^* |
| --- | --- | --- | --- | --- | --- | --- | --- | --- | --- |
| < 0.06500 | 3.226 | 0.1655% to 16.19% | 97.14 | 85.47% to 99.85% | 1.129 | 0.50 | 0.53 | 0.20 | 0.82 |
| < 0.1050 | 22.58 | 11.40% to 39.81% | 91.43 | 77.62% to 97.04% | 2.634 | 0.54 | 0.72 | 0.23 | 0.91 |
| < 0.1350 | 48.39 | 31.97% to 65.16% | 88.57 | 74.05% to 95.46% | 4.234 | 0.63 | 0.81 | 0.30 | 0.94 |
| < 0.1750 | 61.29 | 43.82% to 76.27% | 71.43 | 54.95% to 83.67% | 2.145 | 0.65 | 0.68 | 0.32 | 0.90 |
| < 0.2050 | 77.42 | 60.19% to 88.60% | 48.57 | 32.99% to 64.43% | 1.505 | 0.68 | 0.60 | 0.35 | 0.86 |
| < 0.2150 | 80.65 | 63.72% to 90.81% | 45.71 | 30.47% to 61.81% | 1.486 | 0.70 | 0.60 | 0.37 | 0.86 |
| < 0.2300 | 83.87 | 67.37% to 92.91% | 40 | 25.55% to 56.43% | 1.398 | 0.71 | 0.58 | 0.38 | 0.85 |
| < 0.2450 | 93.55 | 79.28% to 98.85% | 37.14 | 23.17% to 53.66% | 1.488 | 0.85 | 0.60 | 0.59 | 0.86 |
| < 0.2550 | 96.77 | 83.81% to 99.83% | 31.43 | 18.55% to 47.98% | 1.411 | 0.91 | 0.59 | 0.71 | 0.85 |
| < 0.2800 | 96.77 | 83.81% to 99.83% | 28.57 | 16.33% to 45.05% | 1.355 | 0.90 | 0.58 | 0.69 | 0.84 |
| < 0.3150 | 96.77 | 83.81% to 99.83% | 22.86 | 12.07% to 39.02% | 1.254 | 0.88 | 0.56 | 0.64 | 0.83 |
| < 0.3900 | 96.77 | 83.81% to 99.83% | 11.43 | 4.535% to 25.95% | 1.093 | 0.78 | 0.52 | 0.47 | 0.81 |

***^a^*** NPV when treatment-induced survival rate is low (50%)

***^b^*** PPV when treatment-induced survival rate is low (50%)

***^c^*** NPV when treatment-induced survival rate is high (95%)

***^d^*** PPV when treatment-induced survival rate is high (95%)

**Table S9 | Performance of lung [^18^F]FDG uptake reported as %ID/mL at 3 days post-infection (dpi) with CA-09 strain in detecting dampened influenza-associated lung inflammation and predicting survival beyond 6 dpi**. Excerpt of model specificity and sensitivity data obtained from receiving operator characteristic (ROC) analysis in GraphPad Prism v11.0.2. Two sets of negative predictive values (NPV) and positive predictive values (PPV) were calculated based on assumptions of low survival rate (50%) or high survival rate (95%) following therapeutic intervention.

| Cut-off | Sensitivity% | 95% CI | Specificity% | 95% CI | Likelihood ratio | NPV_low_*^a^* | PPV_low_*^b^* | NPV_high_*^c^* | PPV_high_*^d^* |
| --- | --- | --- | --- | --- | --- | --- | --- | --- | --- |
| < 5.065 | 62.5 | 30.57% to 86.32% | 96.55 | 82.82% to 99.82% | 18.13 | 0.72 | 0.95 | 0.39 | 0.99 |
| < 5.170 | 62.5 | 30.57% to 86.32% | 93.1 | 78.04% to 98.77% | 9.063 | 0.71 | 0.90 | 0.38 | 0.97 |
| < 5.255 | 75 | 40.93% to 95.56% | 93.1 | 78.04% to 98.77% | 10.88 | 0.79 | 0.92 | 0.48 | 0.98 |
| < 5.350 | 75 | 40.93% to 95.56% | 89.66 | 73.61% to 96.42% | 7.25 | 0.78 | 0.88 | 0.47 | 0.97 |
| < 5.470 | 87.5 | 52.91% to 99.36% | 89.66 | 73.61% to 96.42% | 8.458 | 0.88 | 0.89 | 0.64 | 0.97 |
| < 5.595 | 87.5 | 52.91% to 99.36% | 86.21 | 69.44% to 94.50% | 6.344 | 0.87 | 0.86 | 0.63 | 0.96 |
| < 5.695 | 87.5 | 52.91% to 99.36% | 82.76 | 65.45% to 92.40% | 5.075 | 0.87 | 0.84 | 0.62 | 0.95 |
| < 5.730 | 87.5 | 52.91% to 99.36% | 79.31 | 61.61% to 90.15% | 4.229 | 0.86 | 0.81 | 0.61 | 0.94 |
| < 5.915 | 87.5 | 52.91% to 99.36% | 75.86 | 57.89% to 87.78% | 3.625 | 0.86 | 0.78 | 0.60 | 0.94 |
| < 6.125 | 87.5 | 52.91% to 99.36% | 72.41 | 54.28% to 85.30% | 3.172 | 0.85 | 0.76 | 0.59 | 0.93 |
| < 6.320 | 87.5 | 52.91% to 99.36% | 65.52 | 47.35% to 80.06% | 2.538 | 0.84 | 0.72 | 0.57 | 0.91 |

***^a^*** NPV when treatment-induced survival rate is low (50%)

***^b^*** PPV when treatment-induced survival rate is low (50%)

***^c^*** NPV when treatment-induced survival rate is high (95%)

***^d^*** PPV when treatment-induced survival rate is high (95%)

**Table S10 | Performance of lung [^18^F]FDG uptake reported as %ID/mL at 5 days post-infection (dpi) with CA-09 strain in detecting dampened influenza-associated lung inflammation and predicting survival beyond 6 dpi**. Excerpt of model specificity and sensitivity data obtained from receiving operator characteristic (ROC) analysis in GraphPad Prism v11.0.2. Two sets of negative predictive values (NPV) and positive predictive values (PPV) were calculated based on assumptions of low survival rate (50%) or high survival rate (95%) following therapeutic intervention.

| Cut-off | Sensitivity% | 95% CI | Specificity% | 95% CI | Likelihood ratio | NPV_low_*^a^* | PPV_low_*^b^* | NPV_high_*^c^* | PPV_high_*^d^* |
| --- | --- | --- | --- | --- | --- | --- | --- | --- | --- |
| < 5.730 | 50 | 21.52% to 78.48% | 100 | 83.89% to 100.0% |  | 0.67 | 1.00 | 0.33 | 1.00 |
| < 5.790 | 62.5 | 30.57% to 86.32% | 100 | 83.89% to 100.0% |  | 0.73 | 1.00 | 0.40 | 1.00 |
| < 5.935 | 62.5 | 30.57% to 86.32% | 95 | 76.39% to 99.74% | 12.5 | 0.72 | 0.93 | 0.39 | 0.98 |
| < 6.095 | 75 | 40.93% to 95.56% | 95 | 76.39% to 99.74% | 15 | 0.79 | 0.94 | 0.49 | 0.98 |
| < 6.355 | 87.5 | 52.91% to 99.36% | 95 | 76.39% to 99.74% | 17.5 | 0.88 | 0.95 | 0.66 | 0.99 |
| < 6.575 | 87.5 | 52.91% to 99.36% | 90 | 69.90% to 98.22% | 8.75 | 0.88 | 0.90 | 0.64 | 0.97 |
| < 6.805 | 87.5 | 52.91% to 99.36% | 85 | 63.96% to 94.76% | 5.833 | 0.87 | 0.85 | 0.63 | 0.96 |
| < 7.035 | 87.5 | 52.91% to 99.36% | 80 | 58.40% to 91.93% | 4.375 | 0.86 | 0.81 | 0.62 | 0.95 |
| < 7.190 | 100 | 67.56% to 100.0% | 80 | 58.40% to 91.93% | 5 | 1.00 | 0.83 | 1.00 | 0.95 |
| < 7.425 | 100 | 67.56% to 100.0% | 75 | 53.13% to 88.81% | 4 | 1.00 | 0.80 | 1.00 | 0.94 |

***^a^*** NPV when treatment-induced survival rate is low (50%)

***^b^*** PPV when treatment-induced survival rate is low (50%)

***^c^*** NPV when treatment-induced survival rate is high (95%)

***^d^*** PPV when treatment-induced survival rate is high (95%)

**Table S11 | Performance of metabolic lung volumes (MLV) at 3 days post-infection (dpi) with CA-09 strain in detecting dampened influenza-associated lung inflammation and predicting survival beyond 6 dpi**. Excerpt of model specificity and sensitivity data obtained from receiving operator characteristic (ROC) analysis in GraphPad Prism v11.0.2. Two sets of negative predictive values (NPV) and positive predictive values (PPV) were calculated based on assumptions of low survival rate (50%) or high survival rate (95%) following therapeutic intervention.

| Cut-off | Sensitivity% | 95% CI | Specificity% | 95% CI | Likelihood ratio | NPV_low_*^a^* | PPV_low_*^b^* | NPV_high_*^c^* | PPV_high_*^d^* |
| --- | --- | --- | --- | --- | --- | --- | --- | --- | --- |
| < 0.02500 | 50 | 8.884% to 91.12% | 100 | 80.64% to 100.0% |  | 0.67 | 1.00 | 0.33 | 1.00 |
| < 0.08500 | 100 | 51.01% to 100.0% | 100 | 80.64% to 100.0% |  | 1.00 | 1.00 | 1.00 | 1.00 |
| < 0.1500 | 100 | 51.01% to 100.0% | 87.5 | 63.98% to 97.78% | 8 | 1.00 | 0.89 | 1.00 | 0.97 |
| < 0.1650 | 100 | 51.01% to 100.0% | 81.25 | 56.99% to 93.41% | 5.333 | 1.00 | 0.84 | 1.00 | 0.96 |
| < 0.1750 | 100 | 51.01% to 100.0% | 50 | 28.00% to 72.00% | 2 | 1.00 | 0.67 | 1.00 | 0.89 |
| < 0.1850 | 100 | 51.01% to 100.0% | 43.75 | 23.10% to 66.82% | 1.778 | 1.00 | 0.64 | 1.00 | 0.88 |
| < 0.1950 | 100 | 51.01% to 100.0% | 31.25 | 14.16% to 55.60% | 1.455 | 1.00 | 0.59 | 1.00 | 0.85 |
| < 0.2050 | 100 | 51.01% to 100.0% | 25 | 10.18% to 49.50% | 1.333 | 1.00 | 0.57 | 1.00 | 0.84 |
| < 0.2150 | 100 | 51.01% to 100.0% | 12.5 | 2.221% to 36.02% | 1.143 | 1.00 | 0.53 | 1.00 | 0.82 |
| < 0.2500 | 100 | 51.01% to 100.0% | 6.25 | 0.3206% to 28.33% | 1.067 | 1.00 | 0.52 | 1.00 | 0.81 |

***^a^*** NPV when treatment-induced survival rate is low (50%)

***^b^*** PPV when treatment-induced survival rate is low (50%)

***^c^*** NPV when treatment-induced survival rate is high (95%)

***^d^*** PPV when treatment-induced survival rate is high (95%)

**Table S12 | Performance of metabolic lung volumes (MLV) at 5 days post-infection (dpi) with CA-09 strain in detecting dampened influenza-associated lung inflammation and predicting survival beyond 6 dpi**. Excerpt of model specificity and sensitivity data obtained from receiving operator characteristic (ROC) analysis in GraphPad Prism v11.0.2. Two sets of negative predictive values (NPV) and positive predictive values (PPV) were calculated based on assumptions of low survival rate (50%) or high survival rate (95%) following therapeutic intervention.

| Cut-off | Sensitivity% | 95% CI | Specificity% | 95% CI | Likelihood ratio | NPV_low_*^a^* | PPV_low_*^b^* | NPV_high_*^c^* | PPV_high_*^d^* |
| --- | --- | --- | --- | --- | --- | --- | --- | --- | --- |
| < 0.07000 | 50 | 8.884% to 91.12% | 100 | 72.25% to 100.0% |  | 0.67 | 1.00 | 0.33 | 1.00 |
| < 0.1000 | 50 | 8.884% to 91.12% | 90 | 59.58% to 99.49% | 5 | 0.64 | 0.83 | 0.31 | 0.95 |
| < 0.1400 | 100 | 51.01% to 100.0% | 90 | 59.58% to 99.49% | 10 | 1.00 | 0.91 | 1.00 | 0.98 |
| < 0.1750 | 100 | 51.01% to 100.0% | 80 | 49.02% to 96.45% | 5 | 1.00 | 0.83 | 1.00 | 0.95 |
| < 0.1950 | 100 | 51.01% to 100.0% | 70 | 39.68% to 89.22% | 3.333 | 1.00 | 0.77 | 1.00 | 0.93 |
| < 0.2100 | 100 | 51.01% to 100.0% | 50 | 23.66% to 76.34% | 2 | 1.00 | 0.67 | 1.00 | 0.89 |
| < 0.2250 | 100 | 51.01% to 100.0% | 40 | 16.82% to 68.73% | 1.667 | 1.00 | 0.63 | 1.00 | 0.87 |
| < 0.2400 | 100 | 51.01% to 100.0% | 30 | 10.78% to 60.32% | 1.429 | 1.00 | 0.59 | 1.00 | 0.85 |
| < 0.2600 | 100 | 51.01% to 100.0% | 10 | 0.5129% to 40.42% | 1.111 | 1.00 | 0.53 | 1.00 | 0.82 |

***^a^*** NPV when treatment-induced survival rate is low (50%)

***^b^*** PPV when treatment-induced survival rate is low (50%)

***^c^*** NPV when treatment-induced survival rate is high (95%)

***^d^*** PPV when treatment-induced survival rate is high (95%)
